# Spatial molecular profiling of mixed invasive ductal-lobular breast cancers reveals heterogeneity in intrinsic molecular subtypes, oncogenic signatures, and mutations

**DOI:** 10.1101/2023.09.09.557013

**Authors:** Osama Shiraz Shah, Azadeh Nasrazadani, Julia Foldi, Jennifer M. Atkinson, Celina G Kleer, Priscilla F. McAuliffe, Tyler J Johnston, Wayne Stallaert, Edaise M da Silva, Pier Selenica, Higinio Dopeso, Fresia Pareja, Diana Mandelker, Britta Weigelt, Jorge S. Reis-Filho, Rohit Bhargava, Peter C. Lucas, Adrian V. Lee, Steffi Oesterreich

**Author notes:** **Corresponding Authors:** Steffi Oesterreich, University of Pittsburgh, 5051 Centre Ave, Pittsburgh, PA 15213. Adrian Lee, University of Pittsburgh, 5051 Centre Ave, Pittsburgh, PA 15213. Current affiliation: AstraZeneca (Jorge S. Reis-Filho). Steffi Oesterreich and Adrian Lee share senior authorship.

## Abstract

Mixed invasive ductal and lobular carcinoma (MDLC) is a rare histologic subtype of breast cancer displaying both E-cadherin positive ductal and E-cadherin negative lobular morphologies within the same tumor, posing challenges with regard to anticipated clinical management. It remains unclear whether these distinct morphologies also have distinct biology and risk of recurrence. Our spatially-resolved transcriptomic, genomic, and single-cell profiling revealed clinically significant differences between ductal and lobular tumor regions including distinct intrinsic subtype heterogeneity (e.g., MDLC with TNBC/basal ductal and ER+/luminal lobular regions), distinct enrichment of senescence/dormancy and oncogenic (ER and MYC) signatures, genetic and epigenetic *CDH1* inactivation in lobular, but not ductal regions, and single-cell ductal and lobular sub-populations with unique oncogenic signatures further highlighting intra-regional heterogeneity. Altogether, we demonstrated that the intra-tumoral morphological/histological heterogeneity within MDLC is underpinned by intrinsic subtype and oncogenic heterogeneity which may result in prognostic uncertainty and therapeutic dilemma.

**Significance:** MDLC displays both ductal and lobular tumor regions. Our multi-omic profiling approach revealed that these morphologically distinct tumor regions harbor distinct intrinsic subtypes and oncogenic features that may cause prognostic uncertainty and therapeutic dilemma. Thus histopathological/molecular profiling of individual tumor regions may guide clinical decision making and benefit patients with MDLC, particularly in the advanced setting where there is increased reliance on next generation sequencing.

## Introduction

Breast cancer is a prevalent disease worldwide^1^ and is the most frequently diagnosed cancer in women in the US, with approximately 300,000 cases per year^2^. The majority (approximately 75%) of cases are classified histologically as invasive breast carcinoma of no special type (IBC-NST), previously described as invasive ductal cancer (IDC). Invasive lobular carcinoma (ILC) is the most common special type accounting for approximately 10-15% of all invasive breast cancer diagnoses^3,4^. Apart from the well-defined histologic and tissue histological differences, there are numerous notable molecular distinctions between ILC and IBC-NST. ILC exhibits enrichment of *CDH1*, *FOXA1*, *TBX3*, *ERBB2* and *PTEN* alterations, while IBC-NST shows a higher frequency of *GATA3*, *MAP3K1*, *TP53* and *MYC* alterations^5^. As a result of *CDH1* alterations, ILC lacks E-cadherin expression, an important cell adhesion molecule^6^, and display cytoplasmic p120 staining while IBC-NST retains membranous expression for both molecules^7^. Clinically, ILC is more common in elderly women, presents as larger tumors and is frequently multicentric and multifocal compared to IBC-NST^8,9^. ILC is usually detected later and leads to higher rates of mastectomy ^10–14^. ILC is also associated with more frequent metastases to urogenital and gastrointestinal tracts as well as to serosal surfaces than IBC-NST^15,16^. Despite presenting with markers of favorable prognosis, such as low grade, low proliferation index, estrogen receptor positive and human epidermal growth factor receptor (HER2) low/negative, patients with ILC have a higher rate of late recurrences and worse long-term prognosis than those with IBC-NST^14,17,18^. Given its distinct prognosis and outcome compared to IBC-NST, ILC is increasingly being recognized as a distinct disease, yet it is still treated similarly to IBC-NST in the clinic due to a lack of specific treatment guidelines^14,19,20^. Ongoing clinical trials focused on patients with ILC may lead to the development of more tailored and effective treatment options in the future^20^.

Apart from ILC, there are numerous other special types of breast cancers such as mixed invasive breast carcinoma. This is an elusive group of breast cancers which the World Health Organization (WHO) defines as tumors presenting with areas of 10-90% of special histology (such as ILC) admixed with at least 10% of IBC-NST^21^. Importantly, this current WHO definition of mixed invasive breast cancers MDLC differs considerably from its previous definition i.e., tumors presenting with areas of 10-49% of the special subtype and at least 50% IBC-NST^22^, indicating the fluid interpretation of this disease. Mixed tumors comprising of both ILC and IBC-NST are the most common (∼5-10% of all breast cancer cases^23–28^) and have different terminologies in the literature including mixed invasive ductal and lobular breast carcinoma (MDLC)^29^ (used throughout this study), mixed invasive ductolobular breast cancer (MIDLC)^30^), invasive ductolobular carcinomas (IDLC)^31^, and more. Despite increasing appreciation of MDLC disease^32,33^, it remains understudied, leading to diagnostic and prognostic uncertainty and clinical management challenges for the patients with this elusive disease. Although there are limited reports focused on MDLC, a few seminal studies have provided insights into its complex pathobiology. Immunohistochemical (IHC) studies have confirmed while that ductal regions of MDLC retain membranous E-cadherin, while lobular regions either lose or display aberrant E-cadherin IHC staining^34–36^. Notably, the current MDLC diagnosis guidelines from WHO only consider the presence of mixed histological patterns and lack recommendations for support from E-cadherin IHC. However, a recent study suggests that the use of E-cadherin IHC can improve diagnostic accuracy of lobular tumor components in ILC and MDLC tumors^37^. Clinical studies have often yielded contradictory findings^24–26,33,34,36,38,39^, but overall, they suggest that MDLC is clinically more similar to pure ILC than IBC-NST^29,40,41^ ^29,40^. These discrepancies may be due to the non-standardized and fluid definition of MDLC disease, limited consideration of its underlying tumor regions and their distinct features and proportions, and the vast intratumoral heterogeneity of this disease^29,33,42,43^. Findings from previous reports point to several subtypes of MDLC tumors based on their histological and E-cadherin staining patterns: 1) intermixed/intermingled type where the E-cadherin positive ductal and E-cadherin negative/aberrant lobular tumor cells appear closely intermixed/intermingled together^31,44^, 2) collision type where the E-cadherin positive ductal and E-cadherin negative/aberrant lobular tumor regions appear in spatially distinct regions which may collide into each other forming intermixed regions^31,45^, and 3) lobular-like IBC-NST type where there are histologically distinct ductal and lobular tumor regions but the overall tumor is predominately E-cadherin positive^46^.

Moreover, molecular studies with profiling of individual tumor regions have been limited. The largest molecular study on MDLC tumors was from the TCGA group^47^, where 88 MDLC were classified into lobular-like and ductal-like groups based on transcriptomic and mutational similarity to ILC and IBC-NST tumors. The lobular-like group comprised of ∼25% of MDLC cases and showed enrichment of mutations in and lower expression of *CDH1* gene. However, these findings may not have any biological relevance as these samples lacked comprehensive annotation and may have been incorrectly defined as MDLC as well as the study lacked individual tumor region analysis. Additional molecular studies have largely focused on analysis of somatic genetic alterations^44,45,48^. While these have provided valuable insights into the tumorigenesis pathways and distinct mutational landscape of ductal and lobular tumor regions^36,49^, comprehensive studies delineating transcriptomic differences between these distinct regions have not been performed.

To fill this knowledge gap, we employed spatially resolved transcriptomic, genomic, and single cell profiling approaches to comprehensively investigate MDLC and highlight its intratumoral heterogeneity and complexity. We delineate the biological and intrinsic subtype differences between its underlying ductal and lobular tumor regions. Notably, our findings highlight that in some MDLC cases distinct tumor regions with distinct biology may pose distinct progression risk. Hence histopathological and molecular profiling MDLC and its individual tumor regions may inform treatment decisions.

## Results

### Patient Specimens and Their Clinicopathologic Characteristics

A panel of three molecular pathologists identified invasive breast cancer cases diagnosed as “mixed invasive ductal and lobular carcinoma” and shortlisted three cases for this pilot study. All three cases were reported as estrogen receptor positive (ER+) and HER2 negative/low and their ages ranged from 46-63 years (Fig. 1A, Supplementary Table S1A). Assessment of H&E and E-cadherin/p120 dual staining confirmed that each case exhibited histomorphic features in line with the latest MDLC definition i.e., admix of E-cadherin negative lobular tumor regions with at least 10-90% of E-cadherin positive ductal tumor regions^21^ (Supplementary Table S1B). Importantly, overall, these cases exhibited collision of spatially distinct ductal and lobular tumor regions^31,45^ along with some regions of intermixing and hence could be broadly defined as MDLC of collision type. These cases were particularly amenable for downstream multi-omic profiling compared to those that exhibited more intermingling or intermixing with close proximity of histologic tumor regions. Clinical diagnostic and treatment timelines of these cases, source of their tumor tissue used in this study, and their histomorphic features are illustrated in Fig. 1A. Both MDLC-2 and MDLC-3 eventually developed recurrent disease resulting in patient death in the case of MDLC-2. As of the last follow-up in 2022, in both MDLC-3 and MDLC-1 cases, the patients were alive with tumors, and in the case of MDLC-1 the patient was free of recurrent disease.

**Figure 1.**
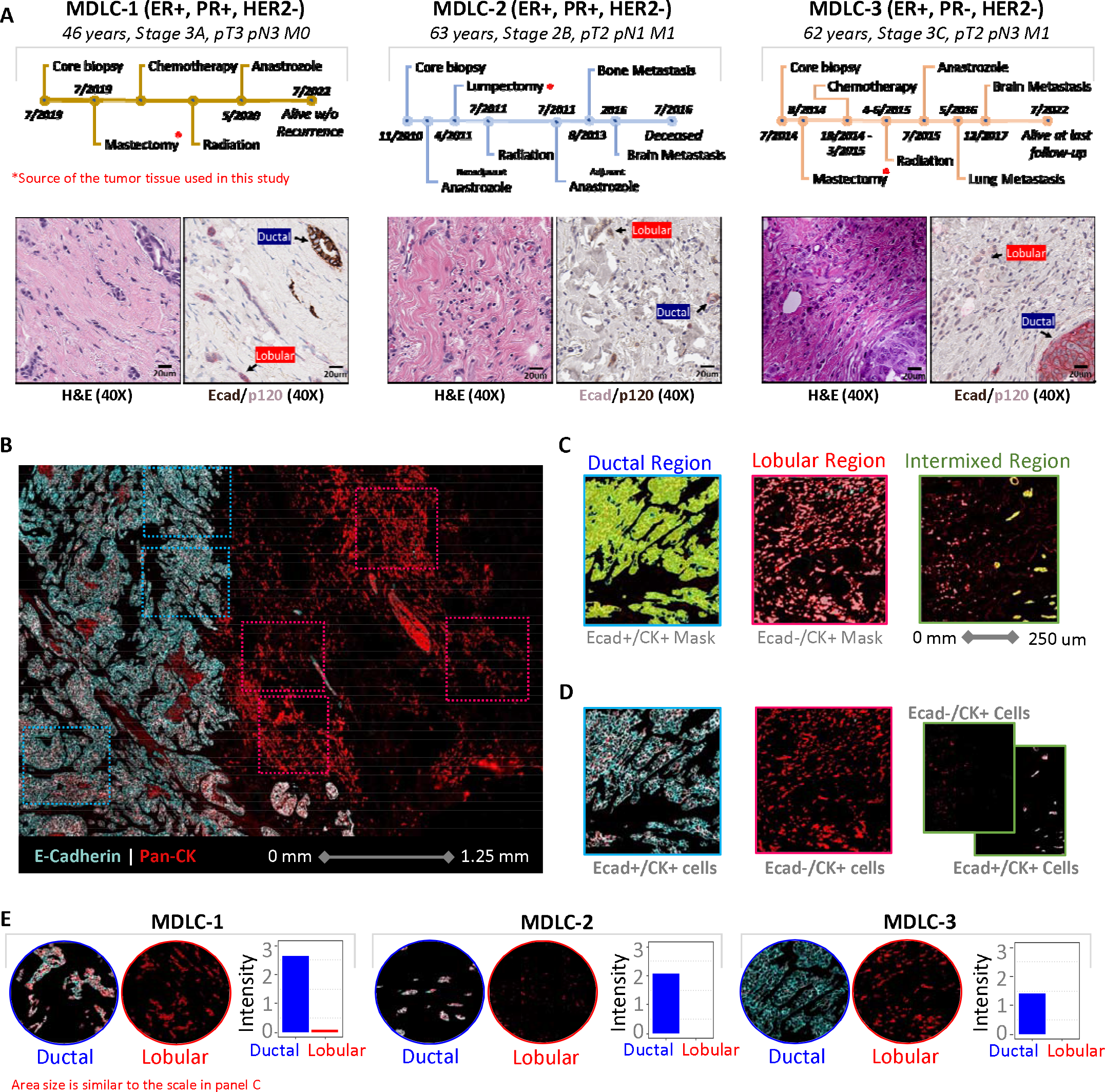
Digital Spatial Profiling of MDLC Ductal and Lobular Tumor Regions. (A) Top panel shows clinical timelines of the studied cases including the time of core biopsies, surgical interventions, treatments, metastasis and last known vital status. The source of the tumor tissue used in this study is highlighted with * in red color. The bottom panels show high (40X) magnification H&E and Ecad/p120 IHC images highlighting histomorphic and E-cadherin features of ductal and lobular regions across each case. (B) E-cadherin and Pan-CK immunofluorescence staining of a representative case (MDLC-3). Selected regions of interests (ROIs) are shown with dotted blue (for ductal) and red (for lobular). (C) Segmentation of ductal and lobular cells using E-cadherin and Pan-CK staining masks. (D) Segmented areas containing only ductal or lobular tumor cells defined using ductal and lobular masks illustrated in panel C. (E) E-cadherin staining intensity in representative ductal and lobular regions profiled across cases.

### Whole-Transcriptome Spatial Profiling of Ductal and Lobular Regions

To better understand the biological heterogeneity and transcriptomic features of ductal and lobular regions, we performed whole-transcriptome digital spatial profiling (DSP) of these three MDLC cases. In total, we analyzed 26 regions of interest (ROIs) with at least 6 ROIs per case. Selected ROIs from a representative case (MDLC-3) are shown in Fig. 1B. We segmented ductal and lobular areas using E-cadherin and Pan-CK staining (Fig. 1C). This enabled profiling of pure ductal and lobular areas from heterogeneous regions that were predominantly of one histology or a mixture of both (Fig. 1D). E-cadherin expression in segmented ductal and lobular regions showed expected results (Fig. 1E). All profiled regions had high RNA sequencing saturation (> 94%) and low noise (Q3 count > Negative probe count). However, three regions had low read count (< 0.3 million) and were excluded from further analysis (Supplementary Fig. S1A-C). Our final DSP dataset is comprised of 13 ductal and 13 lobular regions.

### Transcriptomic Clusters, Intrinsic Molecular Subtypes and Differential Transcriptomic Features

Exploration using top variable genes revealed that regions from each tumor clustered based upon their histology (Fig. 2A; Supplementary Fig. S1D). MDLC-3 ductal regions showed the most distinct mRNA expression compared to all other regions. PAM50 analysis showed both inter and intra-tumor heterogeneity of prognostically relevant intrinsic molecular subtypes (IMS). MDLC-2 ductal regions were a mix of Luminal A and HER2-Enriched subtypes while lobular regions were HER2-Enriched (but stained negative for HER2 IHC – Supplementary Table S1A). MDLC-1 ductal and lobular regions were both of luminal IMS, with the former being Luminal A and the latter being Luminal B. Finally, in line with mRNA expression patterns, MDLC-3 ductal and lobular regions showed stark IMS differences (former Luminal A and later Basal-Like). MDLC-3 ductal region was the only TNBC/Basal-Like tumor (and had low *ESR1*, *PGR* and *ERBB2* mRNA expression – Supplementary Fig. S1E). We further confirmed the TNBC status of MDLC-3 ductal regions using ER and HER2 IHC (Supplementary Table S1A). Overall, our transcriptomic clustering and IMS analysis suggest that ductal and lobular tumor regions within MDLC may have distinct transcriptomes and intrinsic subtypes.

**Figure 2.**
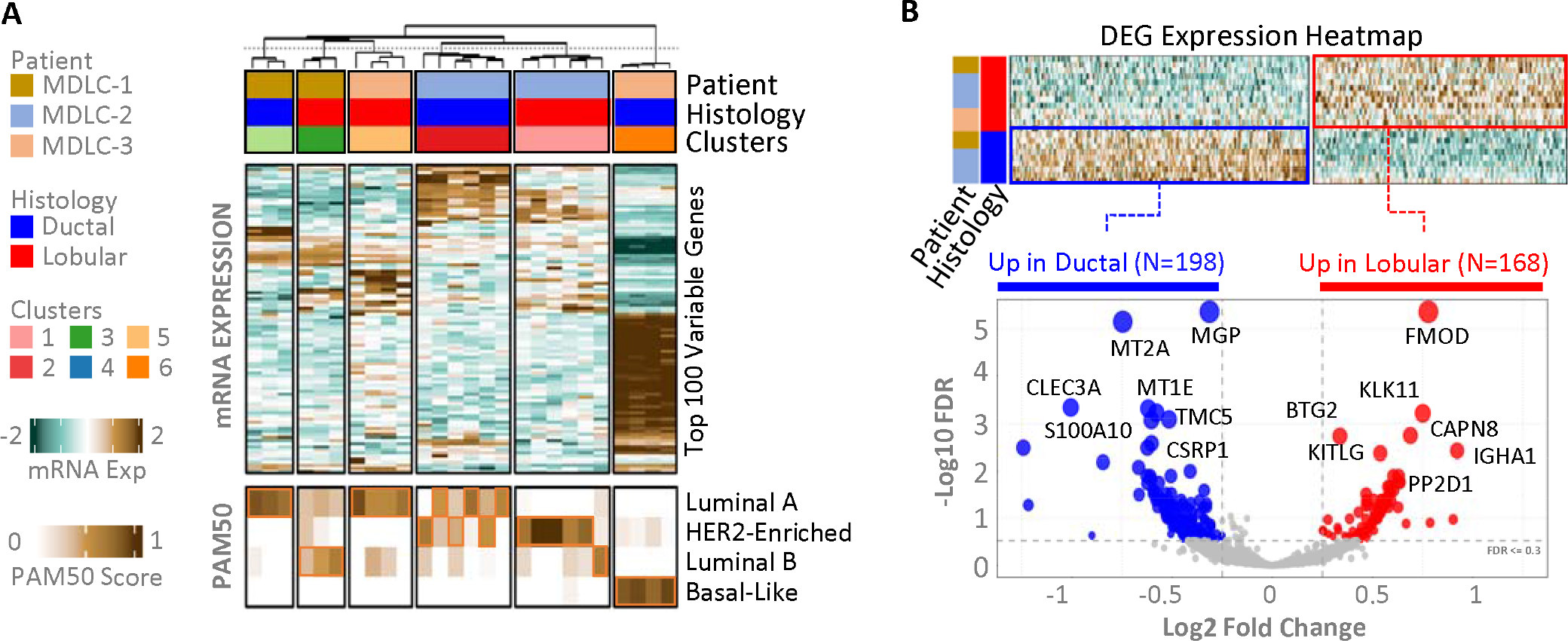
Molecular Subtypes and Differentially Expressed Genes in Ductal and Lobular Tumor Regions. (A) Profiled ductal and lobular tumor regions separated by consensus clusters. All regions are annotated by cases, histology, and consensus clusters. Gene expression heatmap shows top variable genes and followed by PAM50 scores for each subtype across ductal and lobular regions. (B) Differentially expressed genes (DEGs) between ER+ ductal and lobular regions. The top panel shows DEG heatmap, while bottom panel shows volcano plot along with number of significantly upregulated (N=168) and downregulated (N=198) DEGs in ductal vs lobular regions.

To investigate global transcriptomic differences between ductal and lobular regions across all cases, we performed differential gene expression analysis in a pan-patient manner. To avoid biases caused by fundamental differences between ER+ and ER-disease, we excluded MDLC-3 ER-/TNBC ductal regions from this analysis and later downstream analyses (Supplementary Fig. S2C, Supplementary Table S5). In total, we identified 168 significantly up-regulated and 198 significantly down-regulated differentially expressed genes (DEGs) in the lobular compared to ductal regions across all cases (Fig. 2B, Supplementary Table S5). These pan-patient DEGs showed strong overlap with patient specific DEGs (Supplementary Fig. S2A-C) indicating that our pan-patient analysis captured most if not all strong global differences between ductal and lobular tumor regions in each case.

To assess transcriptomic similarity of MDLC ductal and lobular regions to pure counterparts, i.e., IBC-NST (IDC) and ILC, we compared MDLC ductal vs. lobular region DEGs with those from the comparison of TCGA ER+ IBC-NST (N=327) vs ILC (N=113), respectively (Supplementary Fig S2D). We found *KLK11, KLK10, NRAP, SHROOM1, BTG2,* and *PDK4* were upregulated, while *CDH1*, *DCD, SERPINA1*, and *CPB1* were downregulated in both comparisons (i.e., MDLC ductal vs lobular tumor regions and IBC-NST vs ILC tumors). This overlap confirmed the well-known down-regulation of *CDH1* as a marker of the lobular phenotype as well as potential novel markers.

### Differential Enrichment of ER and MYC Signatures & Recurring Biological Themes

To understand the distinct biology of ductal and lobular regions, we scored each DSP region for hallmark signatures^50^. Twenty signatures were differentially enriched between ductal and lobular regions with p-value ≤ 0.05 (Fig. 3A, Supplementary Table S6). Of note, many of these pathways can also be found differentially activated when comparing ER+ IBC-NST and ER+ ILC in TCGA – for example, Kras and TNFA signaling as well as Hypoxia are among significantly enriched pathways in ER+ ILC TCGA samples, whereas Myc targets are enriched in ER+ IBC-NST TCGA samples (Fig. 3B). In line with differential enrichment of Estrogen Response and Myc Target signatures, our transcription factor (TF) signature analysis revealed enrichment of *ESR1/FOXA* TF signatures in lobular and *MYC* TF signatures in ductal regions (Supplementary Fig. S3).

**Figure 3.**
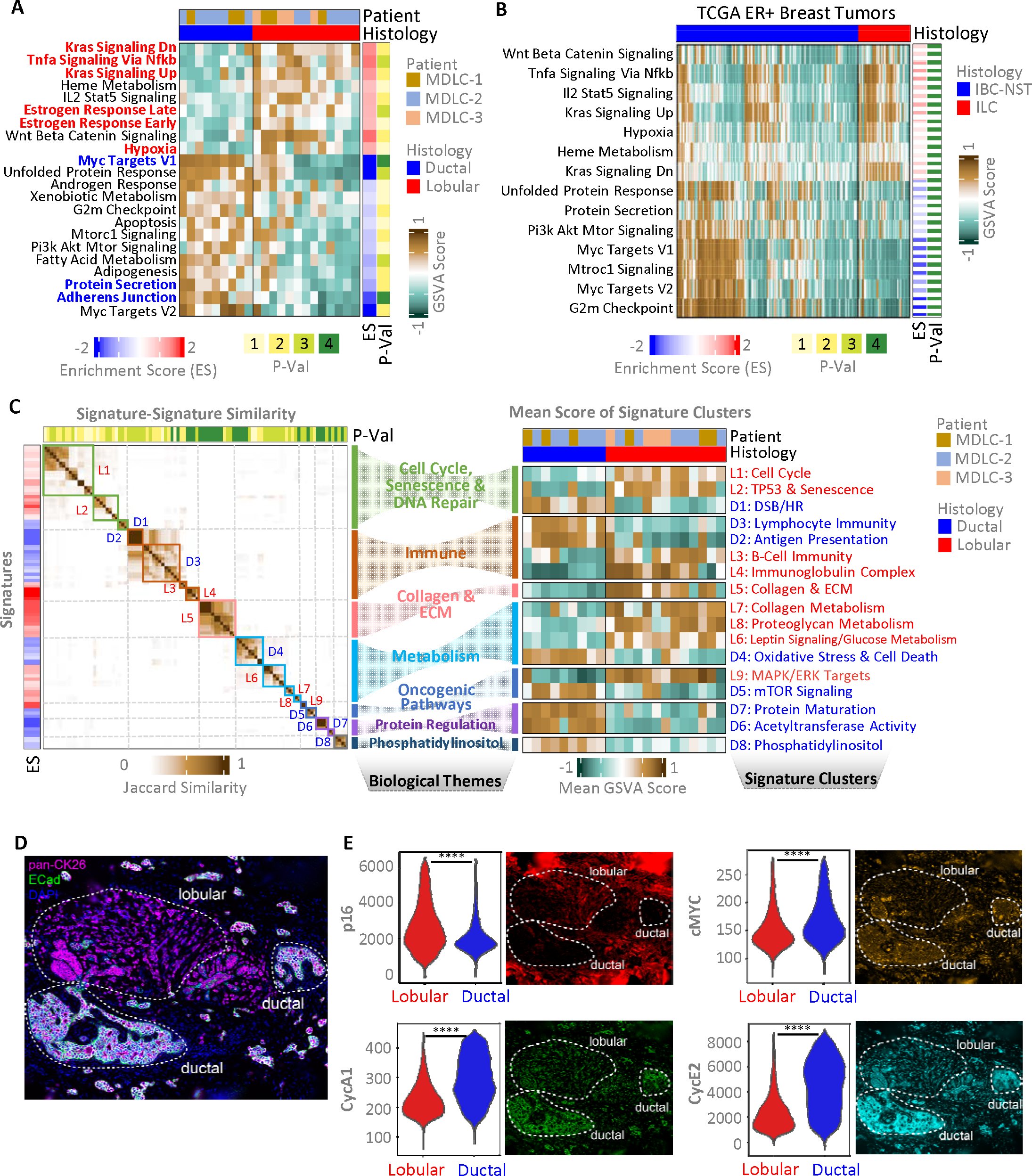
Distinct Hallmark and Biological Signatures in Ductal and Lobular Tumor Regions. (A) Differentially enriched cancer hallmark signatures. The heatmap shows GSVA scores of each differentially enriched pathway in ductal vs lobular tumor regions. Pathways with bold red and blue names are the ones that also were significant in the scRNAseq dataset (Supplementary Fig. S8A). Enrichment score (ES) for a pathway shows mean differences of GSVA score for that pathway in lobular – ductal regions and the −log10 of FDR-adjusted P-value (P-val) shows statistical significance of enrichment based on T test. (P-val score definitions: 4 = P ≤ 0.0001, 3 = P ≤ 0.001, 2 = P ≤ 0.01 and 1 = P ≤ 0.05). (B) The heatmap shows GSVA scores of differentially enriched signatures between TCGA ER+ IBC-NST vs. ILC that overlap with MDLC ductal vs lobular tumor region hallmark signatures in panel A. (C) Recurrent biological themes in ductal and lobular tumor regions. The left panel shows the Jaccard similarity between all signatures. Signatures are grouped based on their Jaccard similarity into signature clusters. Each cluster is associated with a broad biological theme (shown in middle). The right panel shows the mean GSVA score of each signature cluster in ductal and lobular regions. (D) E-cadherin (ECad) and pan-cytokeratin (pan-CK26) staining in representative image of adjacent ductal and lobular tumor regions. (E) Representative immunofluorescence images (right panel) and single-cell quantification (left panel) of cell cycle and senescence biomarkers from all ER+ ductal and lobular regions. Statistical significance is determined using a student’s t-test (****: P ≤ 0.0001). ECM: Extracellular matrix, DSB: Double stranded break, HR: Homologous recombination.

To determine recurring biological themes, we expanded our differential enrichment analysis to include all GO^51^, REACTOME^52^ and KEGG^53^ genesets. Signatures differentially enriched between ductal and lobular regions with p-value ≤ 0.05 were further assessed for robustness against background permutations. Robust signatures were grouped into signature clusters based on similarity of overlapping genes in ductal and lobular regions. All signature clusters were further grouped into broad biological themes based on the underlying biological terms (Fig. 3C, left panel). The mean score of all signature clusters in ductal and lobular regions is shown in Fig. 3C, right panel. Lobular regions were enriched for TP53 & senescence, collagen & extracellular matrix (ECM), MAPK/ERK targets (in line with KRAS signatures shown in Fig. 3A), while ductal regions were enriched in double stranded break & homologous repair (DSB/HR), mTOR signaling (in line with Mtorc1/Mtor signatures shown in Fig. 3A) among other signatures (Supplementary Table S7).

Notably, several genes including *MYC, G3BP1* and *SNRPD1* overlapped between Myc Targets V1 and G2m Checkpoint signatures (both enriched in ductal regions). Both of these signatures had no overlapping genes with DNA damage and repair (D1: DBS/HR) signatures (Supplementary Fig. S4A) indicating that Myc related signatures in ductal regions are associated with proliferation. Importantly, various senescence and cell death related genes (*CDKN1A/2B/2C, TP53, BCL6* among others) overlapped between the L1: Cell cycle and L2: *TP53* & Senescence signatures indicating negative regulation of cell cycle via *TP53* signaling and cell cycle inhibitors (Supplementary Fig. S4B). Key genes associated with these notable and other biological signatures are highlighted in Supplementary Fig. S4C. In summary, we identified various differentially enrichment of various oncogenic and biological signatures between ductal vs. lobular regions including cell cycle (Myc and G2m genes in ductal, and TP53 and cell cycle arrest genes in lobular), immune, collagen & ECM and metabolism among others.

To further investigate some of the notable distinctions in gene expression signatures between ductal and lobular tumor regions, we performed multiplexed immunofluorescence to obtain single-cell proteomic measurements of key cell cycle and senescence effectors in ductal and lobular tumor cells, identified by staining for pan-cytokeratin 26 and E-cadherin (Fig. 3D, Supplementary Table S12). Individual cells within ductal regions exhibited significantly higher expression of proliferative markers including cMYC, cyclin E2, and cyclin A1, while lobular regions had significantly higher expression of the senescence biomarker p16 (Fig. 3E). These findings confirmed the activation of Myc signaling and proliferative pathways in ductal, and activation of cell cycle arrest/senescence in lobular tumor regions.

### Genomic Landscape and Clinically Relevant Genomic Alterations

To explore the genomic landscape of ductal and lobular regions, we employed the FDA-approved targeted MSK-IMPACT panel. We micro-dissected the ductal and lobular tumor regions from FFPE blocks of each case and sequenced them at a median depth of 416x (range 287–563) (Supplementary Fig. S5A). In all cases, at least one pathogenic mutation was detected in known breast cancer driver genes, including *CDH1, PIK3CA, TP53, GATA3, KMT2C, MAPK31, ARID1A,* and *CBFB*, in both ductal and lobular regions, as depicted in Fig. 4A. Furthermore, all pathogenic mutations had a high cancer cell fraction (Supplementary Fig. S5B).

**Figure 4.**
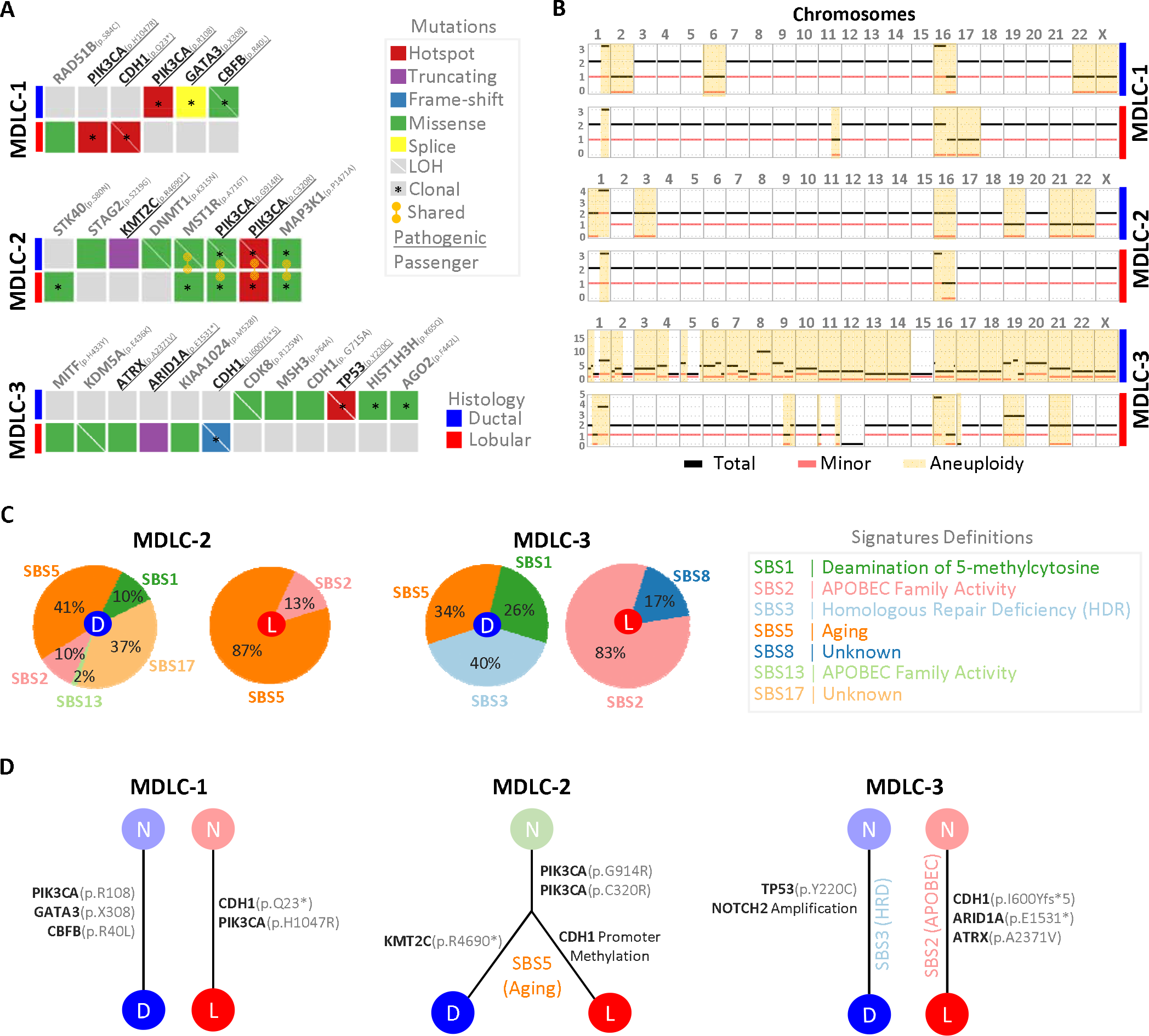
Overview of Genomic Alterations in Ductal and Lobular Tumor Regions. (A) Somatic mutations identified using MSK-IMPACT panel in ductal and lobular components across samples. Gene names are followed by amino acid change due to mutation. Genes in black font had somatic mutations in breast cancer driver genes while those in gray color had passenger/non-driver mutations. (B) Somatic copy number changes in ductal and lobular components across samples. For each case, genome-wide ductal and lobular copy number profiles are shown. Black line shows total copy number, red line shows minor allele copy number and off-white regions show aneuploidy (change in minor allele copy number including LOH). (C) Mutational signatures in MDLC-2 and MDCL-3 ductal and lobular tumor regions. Pie charts show percentage similarity of mutational patterns of each tumor component with established mutational patterns (associated with different means/definition). (D) Predicted disease evolution. Tumors with overlapping mutations were linked to common root node indicating common progenitor or precursor neoplasm (N). Those without common root node likely indicate evolution from independent precursors. Key mutations and events (*CDH1* promoter methylation and amplifications) are shown on the branches.

Loss of E-cadherin is an established driver of lobular tumorigenesis^54^. Typically, it results from biallelic *CDH1* inactivation, through a combination of truncating mutation and loss of heterozygosity (LOH)^28^. All lobular regions exhibited loss of E-cadherin, with both MDLC-1 and MDLC-3 harboring *CDH1* truncating mutations and LOH (Fig. 4B; Fig. 1E; Supplementary Fig. S1E). The lobular region of MDLC-2 did not have *CDH1* genomic alterations but showed epigenetic inactivation via methylation of its promoter. A putative passenger mutation in *CDH1* was also observed in the MDLC-3 ductal region, which had no effect on mRNA or E-cadherin levels (Fig. 1E; Supplementary Fig. S1E). The loss of E-cadherin is consistent with the downstream down-regulation of Adherens Junction signatures in all lobular regions (Fig. 3A). Hotspot *PIK3CA* mutations, which are early oncogenic driver events frequently observed in ER+ BC^55^, were present in all cases except MDLC-3. The MDLC-1 ductal region had mutations in *GATA3* (critical transcriptional binding partner of ERalpha^56^) and *CBFB* (forms complex with *RUNX1* to suppress NOTCH signaling^57^). The MDLC-3 ductal region had loss of *TP53*, an event associated with more aggressive types of breast cancer like basal/TNBC^58^ and consistent with its basal-like intrinsic subtype. MDLC-2 ductal region had truncating mutation in *KMT2C*, an important regulator of ER alpha activity^59^. MDLC-3 lobular region had mutations in *ATRX* and *ARID1A,* two chromatin remodeling and DNA repair genes^60^. In addition, while MDLC-1 and MDLC-3 ductal and lobular regions did not share any somatic clonal mutations, MDLC-2 ductal and lobular regions shared several somatic clonal mutations in *PIK3CA* and *MAPK31* genes indicating shared tumor origin.

To investigate the copy number (CN) landscape of ductal and lobular regions, we performed CN analysis using FACETS^61^ (Fig. 4B). All regions exhibited breast cancer characteristic chromosome 1q gains and 16q losses^58^. Overall, ductal tumor regions showed more frequent aneuploidy events than lobular tumor regions. MDLC-3 ductal region showed the most aberrant CN profile consistent with its basal/TNBC etiology^58^. Moreover, chromosome 22 aneuploidy was only observed in ductal tumor regions. Notably, both histologic regions of MDLC-3 and ductal region of MDLC-2 had *NOTCH2* gains or amplifications (Supplementary Fig. S5C).

### Mutational Processes Driving Tumor Evolution

To determine the mutational processes underlying the evolution of ductal and lobular tumor regions within MDLC, we performed mutational signature analysis of all cases (Fig. 4C). Both histologic regions in MDLC-2 were associated with an “aging” signature. In MDLC-3, ductal region was associated with homologous repair deficiency (HRD) while the lobular regions with APOBEC signature. Tumors with an APOBEC signatures have a higher frequency of mutations in chromatin remodeling and DNA repair genes^62^, as seen in MDLC-3 lobular region. We did not identify any signatures for MDLC-1 ductal and lobular regions. The putative evolution of each MDLC case and its histologic region based upon the mutations, *CDH1* alterations and mutational signatures is shown in Fig. 4D. Notably, the MDLC-2 ductal and lobular regions not only shared clonal pathogenic (*PIK3CA*) and passenger (*MST1R* and *MAP3K1*) mutations but also the “aging” mutational signature. Loss of function *CDH1* alterations were hallmark of all lobular regions across cases.

### Single-Cell RNA-Sequencing and Identification of Ductal and Lobular Populations

To explore the heterogeneity of MDLC at a single-cell level, we performed scRNAseq sequencing of a fresh tumor specimen from MDLC-1. The resulting scRNAseq comprised of 4,671 cells and had sequencing saturation, reads in cell and median genes per cell as shown in Supplementary Fig. S6A. Cell-type annotation based on canonical markers (see details in Supplementary Methods) revealed various tumor and non-tumor cell types (Supplementary Fig. S7B). scRNAseq inferred copy number revealed that only the epithelial tumor cells had aneuploid copy number status further confirming their nature (Supplementary Fig. S7C). For the downstream analyses, we only focused on the tumor cell cluster. Clustering of this revealed two transcriptionally distinct tumor cell populations with similar cell cycle profiles; however, we were unable to annotate them as either ductal or lobular tumors using *CDH1* mRNA expression (Fig. 5A). Notably, while the MDLC-1 lobular regions showed loss of E-cadherin protein (Fig. 1E), the *CDH1* p.Q23* truncating mutation in these regions (Fig. 4A) didn’t alter *CDH1* mRNA expression (Supplementary Fig. S1E). Previous studies suggest that some protein truncating mutations can escape non-sense mediated decay and thereby maintain gene expression levels but lack a functional protein^63–65^. Indeed, similar effects of Q23 mutations were seen in the TCGA cohort, where *CDH1* mRNA levels in Q23* mutant tumors were similar to those in *CDH1* WT tumors, while the E-cadherin protein levels in Q23* mutant tumors were similar to the tumors with all other *CDH1* truncating mutations (Supplementary Fig. S6). ^63–65^

**Figure 5:**
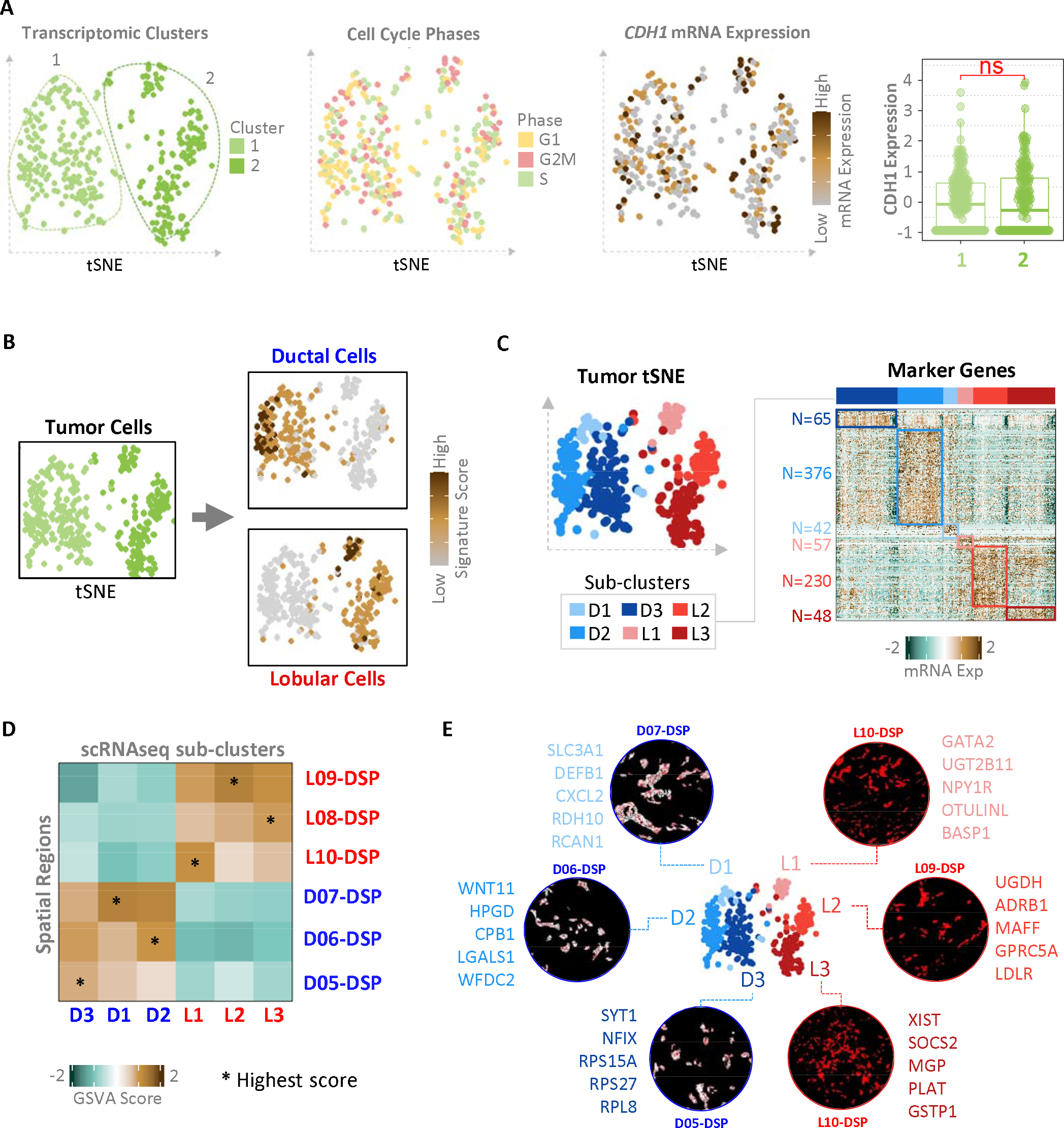
Single Cell Heterogeneity of Ductal and Lobular Tumor Populations. (A) Tumor populations in MDLC-1 scRNAseq dataset (left) and assessment of CDH1 expression in MDLC-1 scRNAseq dataset (middle and right plot). Statistical significance was evaluated using the T test. (B) Deconvolution of scRNAseq using DSP derived ductal and lobular signatures. GSVA scores for ductal and lobular signatures are shown on the right panel. (C) Sub-clustering and identification of ductal and lobular sub-populations (left) with unique marker genes (right). (D) Scoring spatial regions using GSVA of sub-cluster marker genes. (E) Mapping of single cell populations to spatial regions based on GSVA scores in panel D.

To identify ductal and lobular single populations, we derived ductal and lobular transcriptome signatures (Supplementary Table S5) from differential gene expression analysis of the MDLC-1 DSP ductal vs lobular tumor regions and used these signatures to score each single cell in MDLC-1 scRNAseq dataset. Using this deconvolution approach, we successfully annotated ductal and lobular tumor populations in the scRNAseq dataset (Fig. 5B). scRNAseq inferred copy number profiles of ductal and lobular cells revealed various breast cancer specific aberrant/aneuploid events in chromosomes 1, 8, 11, 16, 18 among others (Supplementary Fig. S7E). Overall, the copy number profiles of ductal tumor cells were more aberrant than those of lobular tumor cells. Ductal and lobular tumor regions DEGs in scRNAseq and DSP datasets showed significant overlap (Supplementary Fig. S7D,F). Differential enrichment of several hallmark signatures and recurring biological themes in ductal and lobular regions from DSP data was recapitulated in our scRNAseq data (Supplementary Fig. S8A,C). Notably, we saw enrichment of Estrogen and senescence signatures in lobular and Myc signature in ductal tumor cells in both DSP and scRNAseq datasets.

### Single-Cell Heterogeneity and Mapping of Single-Cells to Spatial Regions

To investigate single-cell heterogeneity of ductal and lobular populations, we increased resolution of clustering to identify sub-clusters. Each sub-cluster had unique marker genes (Fig. 5C) and several of these also showed differential enrichment of hallmark signatures and recurring biological theme clusters (Supplementary Fig. S8B,D), indicating intra-tumor heterogeneity within ductal and lobular tumor populations. To investigate whether single-cell transcriptomic heterogeneity was associated with spatial heterogeneity, we mapped scRNAseq sub-clusters to DSP regions. To assess similarity between each region and sub-cluster, we scored all regions for sub-cluster marker genesets using GSVA (Fig. 5D). Each region was mapped to a sub-cluster for which it had most similarity (i.e., highest GSVA score) (Fig. 5E). The top five marker genes of each scRNAseq sub-cluster (Supplementary Fig. S7G) are shown beside each sub-cluster and spatial region link. These findings perhaps suggest that the single-cells in each sub-cluster came from a spatially distinct niche in the tumor.

## Discussion

Despite an increasing incidence of MDLC^32,33^, this special subtype of breast cancer remains poorly understood, with limited knowledge about its overall biology and that of its underlying ductal and lobular tumor regions, as well as their clinical implications.

Spatially resolved and single-cell profiling techniques have revolutionized our understanding of physiological and pathological processes and tissue architecture^66^. Efforts such as Human Cell Atlas^67^ have successfully uncovered the major cell types and their spatial organization in both normal breast tissue^68,69^ and breast cancer^70^. Spatial and single-cell profiling of primary breast cancer subtypes and metastatic disease have been instrumental in identifying novel molecular features and their prognostic value^71–75^. However, to date genome-wide spatial and single-cell investigation of MDLC has not been reported. To address this gap, we employed spatially resolved transcriptomics, genomics, and single-cell profiling to investigate the biology of individual tumor regions within MDLC of collision type. Our study confirmed various previously reported observations of genetic inactivation of *CDH1* and concomitant loss of E-cadherin in lobular regions^31,34–36^ and the presence of distinct driver alterations in individual tumor regions^31,36^. We also uncovered novel findings that highlight the vast heterogeneity within MDLC, several of which may be indicative of distinct prognosis, hence highlighting the significance of reporting this heterogeneity in clinical diagnosis. Ductal and lobular tumor regions had distinct biological signatures, in particular cell cycle arrest/senescence, ER and *MYC* signaling, and intrinsic molecular subtypes (IMS). The IMS distinctions were most prominent in the case of MDLC-3, which was clinically reported as ER+ tumor but molecular profiling revealed that the underlying ductal region was ER-/TNBC with basal-like IMS while the lobular region was ER+ with luminal-A IMS. Notably, both ER-status and basal-like IMS are associated with worse prognosis and outcome than ER+ and luminal IMS^76,77^. In this case, the patient had brain and lung metastases, both of which were also ER-, suggesting progression of the primary ER-ductal tumor. Another notable finding was enrichment of cell cycle arrest/senescence and ER signaling signatures in lobular while *MYC* signatures in ductal tumor regions. Multiplex immunofluorescence staining of various effector proteins confirmed that Myc and Cell proliferation signaling were active in ductal tumor cells, while cell cycle arrest/senescence signaling was active in lobular regions. Further analysis at single-cell RNA level revealed that ER signatures were enriched in only one lobular subcluster (L2), highlighting molecular heterogeneity even within individual histologically distinct tumor regions. Majority of ILC are strongly ER+ and respond well to endocrine therapies, while they are mostly less proliferative, responding poorly to chemotherapy^14,20,28,78,79^. Whether this is also true for lobular regions within MDLC is an important question and must be further investigated. Perhaps, future trials can study whether tumors with predominantly ductal vs. lobular MDLC respond differently to endocrine and chemotherapies.

Development of robust mutational signature analysis tools such as Signature Multivariate Analysis (SigMa) have allowed accurate identification of mutational signatures associated with such HRD and APOBEC defects from targeted sequencing panels such as MSK-impact^80–84^. Utilization of SigMa algorithm in our study revealed that individual tumors regions with distinct alterations may share common (Aging in MDLC-2) or distinct (APOBEC or HRD in MDLC-3) mutational signatures. Collectively, our results indicate ductal and lobular regions in MDLC-2 have shared origin, while MDLC-3 is a clear collision of two independent tumors. The etiology of MDLC-1 is questionable – while there are no shared mutations, we note that this interpretation is based upon a small panel of genes measured by the MSK IMPACT panel with strict filtering. We cannot exclude the possibility of shared mutations in other cancer-related genes not covered by MSK-impact panel. Larger efforts using whole genome profiling of more MDLC tumors are needed to comprehensively investigate the mutational signatures and origins of ductal and lobular tumor regions in mixed tumors.

The presence of distinct clinically actionable alterations in individual tumor regions within MDLC is of clinical relevance and has been previously reported MDLC^45^. Tumor regions with distinct alterations can complicate clinical management. Alterations in components of ER signaling (such as *GATA3, KMT2C*) as well as Notch signaling (*CBFB*, *NOTCH2*) can lead to endocrine resistance^85–87^, while alterations in DNA repair response genes (*ARID1A*, *ATRX*) may confer increased sensitivity to chemotherapy and other DNA repair pathway targeting drugs including PARP inhibitors^88,89^. Future studies will need to address the important question whether treatment decisions for patients with MDLC benefits from considering the distinct alterations in individual tumor regions.

For a long time, it was unclear whether ductal and lobular regions within MDLC share any molecular similarity with their pure counterparts (i.e., ILC and IBC-NST tumors). To fill in the gap, in our study, we compared the molecular features of individual tumor regions within MDLC with their pure counterparts (i.e., ILC and IBC-NST tumors) in the TCGA cohort^28^ and discovered a set of shared markers of lobular phenotype. There was some but limited overlap of the differentially expressed genes between ductal and lobular tumor regions in MDLC cases with differential gene expression in ER+ ILC vs ER+ IBC-NST in the TCGA cohort. This could be due to technical reasons such as bulk sequencing of frozen tumor samples (in TCGA) vs. spatial sequencing of epithelial tumor regions from FFPE tissues (in this study). Furthermore, this also could also be reflective of true biology i.e., unique gene expression patterns in MDLC tumor regions compared to pure ILC and IBC-NST tumors. As expected, one of overlapping genes that was downregulated in both MDLC lobular tumor regions and ILC tumors was *CDH1*, an important driver of lobular tumorigenesis^54^. ^2854^This primarily occurs via genetic inactivation of *CDH1* in ILC tumors^28^, similar to what was observed in the case of lobular regions in MDLC-1 and MDLC-3. However, in the lobular region of MDLC-2, *CDH1* promoter methylation was observed. *CDH1* promoter methylation is also associated with epithelial-to-mesenchymal transition (EMT), however, EMT is not a typical feature of lobular tumors and is primarily associated more complex subtypes of breast cancers such as metaplastic and claudin-low subtypes^90–93^. EMT markers^90^ (Supplementary Fig. S1F) were not consistently upregulated in MDLC-2 lobular region hence ruling out EMT-associated *CDH1* promoter methylation. This is an intriguing observation, suggesting that the pathogenesis of lobular regions within MDLC may occur via both genetic and epigenetic inactivation of *CDH1,* unlike ILC where genetic inactivation of *CDH1* is predominant^6,31,94–96^. A recent study revealed the synthetic lethality in E-cadherin deficient breast cancers with inhibition of *ROS1*, a tyrosine kinase^97^, and trials are ongoing testing efficacy of *ROS1* inhibitors in ER+ ILC disease. Future studies could assess *ROS1* inhibitors or other drugs in treating patients with predominantly lobular MDLC disease. Additional studies are also warranted to investigate whether E-cadherin deficient tumors with distinct etiology, such as *CDH1* epigenetic inactivation, have distinct biology potentially associated with differential treatment response.

Our study focused primarily on profiling MDLC of collision type as these were the most amenable for our molecular profiling approaches i.e., we could clearly annotate ductal and lobular tumor regions based on histological and E-cadherin staining patterns and cleanly capture RNA/DNA from these individual regions. Notably, we did attempt to profile ductal and lobular tumor cells separately from intermixed regions identified in MDLC-2, however, the resulting data was of poor quality due to low tumor cellularity. Although our study presents a unique and novel analysis of MDLC of collision type, it has some limitations, including a small sample size and lack of analysis of other subtypes (such as lobular-like IBC-NST^46^ and intermixed/intermingled MDLC^31,44^) and tumor microenvironment of individual tumor regions within MDLC. Future studies, using rapidly improving spatial sequencing technologies with increased single cell capabilities will be necessary to validate and expand our observations in larger and more diverse cohorts of MDLC. Generating a comprehensive MDLC cohort for future studies may not be an easy task, given the challenges associated with the diagnosis of this mixed entity. However, the use of E-cadherin IHC in histological assessments and evaluation by a panel of pathologists can ensure the generation of a high-confidence and well-annotated MDLC cohort for future studies. Moreover, the application of digital pathology and artificial intelligence will make this more feasible^98^. These tools will not only improve diagnostic accuracy^99^ but will also allow scalable annotation of ductal and lobular tumor regions across heterogenous tumor samples^100^.

In summary, our study provides new insights into the molecular heterogeneity of MDLC and its underlying ductal and lobular tumor regions. This is the first report on genome-wide spatially resolved and single-cell molecular profiling of MDLC. We discovered distinct biological signatures and intrinsic molecular subtypes between these regions that may have important implications for patient prognosis and clinical management. Our findings also highlight the need for comprehensive histopathological and molecular profiling of individual tumor regions within MDLC to better understand its progression and response to cytotoxic, endocrine, and other targeted therapies.

## Methods

### Sample Collection

Informed patient consent was obtained for tissue collection, and institutional review board approval was obtained from the University of Pittsburgh prior to initiation of the study. Patients diagnosed with “mixed invasive ductal and lobular” were identified from the UPMC Breast Cancer Registry and Formalin-Fixed Paraffin-Embedded (FFPE) blocks requested from Pitt Biospecimen Core. Four-um-thick FFPE sections were used for H&E, ER-alpha and E-cadherin IHC staining. A panel of pathologists reviewed the H&E and E-cadherin staining to confirm MDLC diagnosis in line with recent WHO guidelines^21^ (10-90% of ILC and at least 10% IBC-NST) and shortlisted MDLC cases amenable for multi-omic profiling i.e., those with good separation of ductal and lobular tumor regions to avoid spill over and contamination of individual ductal and lobular tumor regions by neighboring lobular and ductal tumor cells, respectively. Hence, we selected three collision type MDLC cases that showed both clear separation of ductal and lobular tumor regions along with some intermixed/intermingled regions. These included cases MDLC-1, MDLC-2 & MDLC-3 (clinical features and pathology review notes are shown in Supplementary Table S1A & S1B). All cases underwent NanoString digital spatial, MSK-IMPACT mutation and *CDH1* promoter methylation profiling. Fresh tumor was only available for MDLC-1 and was utilized for single-cell RNAseq. See details in Supplementary Methods.

### Multiomic Profiling

Digital spatial profiling (DSP) was performed using NanoString GeoMX whole transcriptome atlas platform. Mutation profiling was performed using MSK-IMPACT targeted NGS panel. *CDH1* promoter methylation was assessed using ddPCR system. Finally, single cell RNA sequencing was performed using 10X genomics platform based on 3’end chemistry v3. Please see supplementary methods for details.

### Multiplexed immunofluorescence

Tissue sections were prepared and iteratively stained as described previously^101^. Briefly, tissue sections were baked overnight at 601C, deparaffinized with xylene, rehydrated by decreasing ethanol washes, followed by proprietary antigen retrieval optimized for multiple antigens (Leica Biosystems). Tissues were incubated in 0.3% Triton X-100 in PBS for 10 min followed by blocking in 10% (w/v) donkey serum and 3% (w/v) bovine serum albumin (BSA) in PBS for 1 h at room temperature. Primary antibodies were diluted in blocking solution and incubated for 1 h at room temperature (anti-E-cadherin, Cell Signaling Technology, 96743; anti-pan-cytokeratin, Millipore Sigma, C5992; anti-c-Myc Alexa Fluor 647, Abcam, ab190560; anti-p16 Alexa Fluor 647, Abcam, ab192054; anti-cyclin E2 Alexa Fluor 488, Abcam, ab207336; anti-cyclin A1, R&D Systems, MAB7046; anti-p21, R&D Systems, AF1047; anti-phospho-H2AX Alexa Fluor 555, Abcam, ab206900). In-house primary antibody fluorescent conjugates were generated using the Alexa Fluor 647 labelling kit (Invitrogen, A20186) or the Alexa Fluor 555 labelling kit (Invitrogen, A20187). Nuclear labelling was performed using Hoechst (2 **u**g/ml, Invitrogen, H3570) for 10 min at room temperature. Imaging was performed in 50% glycerol (v/v) in PBS. After imaging, dye inactivation was performed in an alkali solution containing H2O2 for 15 min with agitation followed by a PBS wash. Samples were reimaged following dye inactivation to measure residual fluorescence. Additional rounds of antibody staining and imaging were performed as described above, starting with primary antibody incubation. Image acquisition, flat-field correction, autofluorescence removal, and registration were performed using the Cell DIVE Imager (Leica Biosystems). Single-cell segmentation was performed using Cellpose^102^ and intensity measurements were extracted using scikit-image^103^. Measurements greater than 1.5x the interquartile range were identified as outliers and removed from analysis.

### Figures & Illustrations

DSP spatial images were exported from GeoMx® Data Analysis Suite (DSPDA) and used directly in the manuscript without graphical modifications. Most bioinformatic analysis visualizations were generated using R. These visualizations were aesthetically improved in power point to produce the final figures. Illustrations were prepared using BioRender.com.

## Data Availability

Data and code generated in this study is shared in the Supplementary Data files, Mendeley (https://data.mendeley.com/datasets/btv7g7n9ys/1) and GitHub (https://github.com/osamashiraz/MDLC_Spatial_Analysis_2023).

## Authors’ Contributions

**O.S. Shah:** Conceptualization, data curation, software, formal analysis, investigation, visualization, methodology, project administration, writing–original draft, writing–review and editing. **A. Nasrazadani:** resources, funding acquisition, writing-review and editing. **J.M. Atkinson:** resources, supervision, project administration, writing–review and editing. **J. Foldi:** resources, investigation, data curation, writing–review and editing. **C.G. Kleer:** investigation, data curation, writing–review and editing. **P.F. McAuliffe:** investigation, resources, data curation, writing–review and editing. **T.J. Johnston** : investigation, resources, data curation, software, formal analysis, writing–review and editing. **W. Stallaert** : investigation, resources, data curation, software, formal analysis, writing–review and editing. **E.M. da Silva:** investigation, resources, data curation, software, formal analysis, writing–review and editing. **P. Selenica:** investigation, resources, data curation, software, formal analysis, writing–review and editing. **H. Dopeso:** investigation, resources, data curation, writing–review and editing. **F. Pareja:** investigation, resources, data curation, writing–review and editing. **D. Mandelker:** investigation, resources, data curation, writing–review and editing. **B. Weigelt:** conceptualization, investigation, methodology, resources, writing–review and editing. **J.S. Reis-Filho:** conceptualization, investigation, methodology, resources, writing–review and editing. **R. Bhargava:** conceptualization, investigation, resources, data curation, methodology, data curation, writing–review and editing. **P.C. Lucas:** conceptualization, investigation, resources, data curation, methodology, data curation, writing–review and editing. **A.V. Lee:** conceptualization, resources, methodology, supervision, funding acquisition, project administration, writing–review and editing. **S. Oesterreich:** conceptualization, resources, methodology, supervision, funding acquisition, project administration, writing–review and editing.

## Acknowledgments

This work was primarily supported by the Dynami Foundation (Flora Migyanka) and BCRF grants to **S. Oesterreich** and in part by various other grants including the ASCO Gianni Bonadonna Breast Cancer Research Fellowship grant to **A. Nasrazadani** and Conquer Cancer ASCO Young Investigator Award to **J. Foldi**. We would also like to thank NanoString DSP Technology Access Program for providing early access to their whole transcriptome spatial technology, UPMC Hillman Cancer Center and Tissue and Research Pathology/Pitt Biospecimen Core shared resources for providing access to valuable clinical specimens and technical support (award P30CA047904), University of Pittsburgh Center for Research Computing, RRID:SCR_022735 (HTC cluster, NIH Award S10OD028483), Pitt Genomics Research Core and UPMC Genome Center for technical support in preparing scRNAseq libraries and sequencing, and Ms. Morgan DeBerry, HTL(ASCP)CM, QIHCCM for excellent technical assistance in tissue sectioning, staining and other logistics. Research reported in this publication was partly funded by an NIH/NCI Cancer Center Support Grant (P30CA008748, MSKCC). BW, FP and JSR-F were funded in part by an NIH/NCI P50 CA247749 01 and Breast Cancer Research Foundation grants.

## Supplementary Figure Legends

**Supplementary Figure S1:**
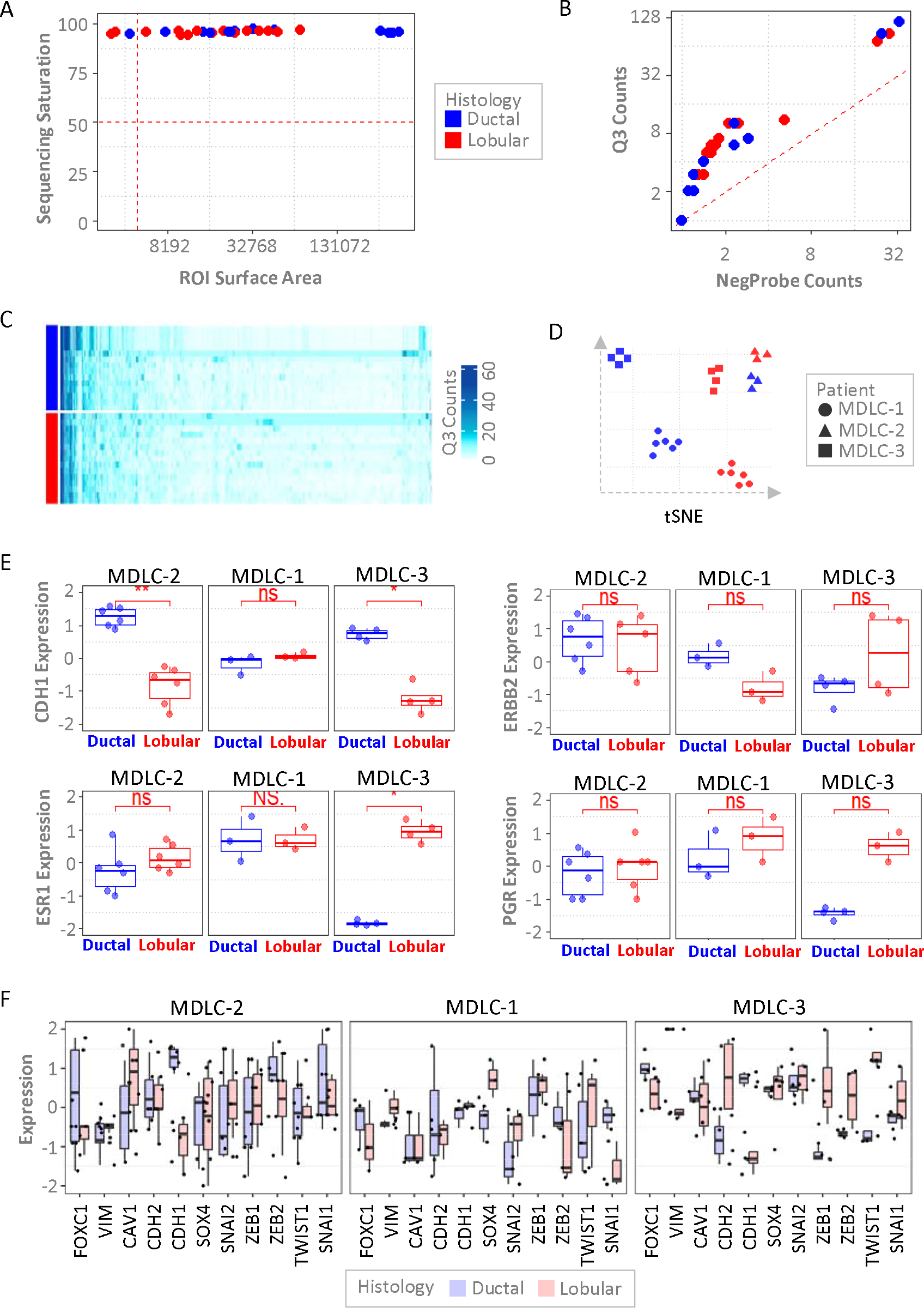
Digital Spatial Profiling RNAseq Dataset Quality Control. (A) Scatter plot showing sequencing saturation (y-axis) and area of interest (AOI) surface area (mm2) across cases. Dashed red lines show the quality control (QC) cutoffs (sequencing saturation >50% and area > 6000mm^2^) for selecting high quality samples. (B) Overview of signal (Q3-normalized counts) to noise (negative probe counts). Dashed red line shows cutoff i.e., signal > noise. (C) Evaluation of Q3 normalized counts (for 1000 randomly sampled genes) across profiled regions to identify samples with low quality data showing count artifacts. (D) tSNE plot showing clustering of profiled samples that passed QC cutoffs. (E) *CDH1, ESR1, ERBB2* and *PGR* gene expression in ductal vs lobular regions across the three profiled cases. Statistical significance was evaluated using the T test. (F) Expression of EMT marker genes in ductal and lobular tumor regions across each case. EMT marker genes were adapted from McCart Reed et al^34^.

**Supplementary Figure S2:**
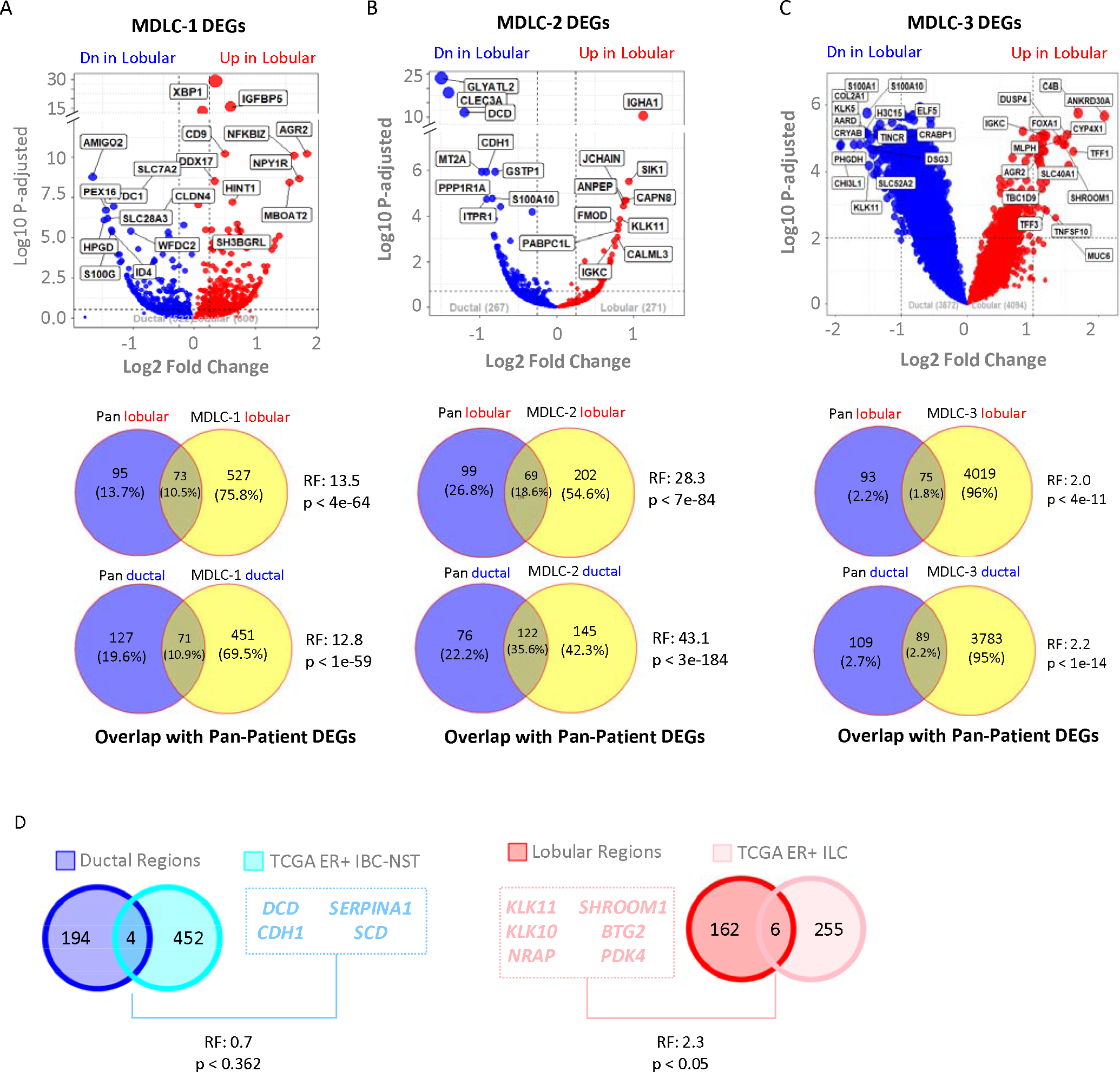
Patient Wise and Pan-Patient MDLC Ductal vs Lobular Differential Analysis. Volcano plots (top) showing per patient or patient-wise differentially expressed genes (DEGs) between ductal vs lobular tumor regions and overlap of these patient-wise DEGs with pan-patient (all patients combined) DEGs for (A) MDLC-1, (B) MDLC-2 and (C) MDLC-3. A RF > 1 and p < 0.01 shows significantly strong overlap. (D) Overlap of MDLC pan-patient ductal vs. lobular DEGs with TCGA ER+ IBC-NST vs ILC DEGs. Statistical significance of overlap between genesets was computed using overlap_stats tool (http://nemates.org/MA/progs/overlap_stats.html). RF: Representation Factor. For details on RF calculation see overlap_stats webpage. Briefly, RF is the number of overlapping genes divided by the expected number of overlapping genes drawn from two independent groups. A RF > 1 indicates more overlap than expected for independent genesets, a RF < 1 indicates less overlap than expected, and a RF = 1 indicates that the two genesets by the number of genes expected for independent groups of genes.

**Supplementary Figure S3:**
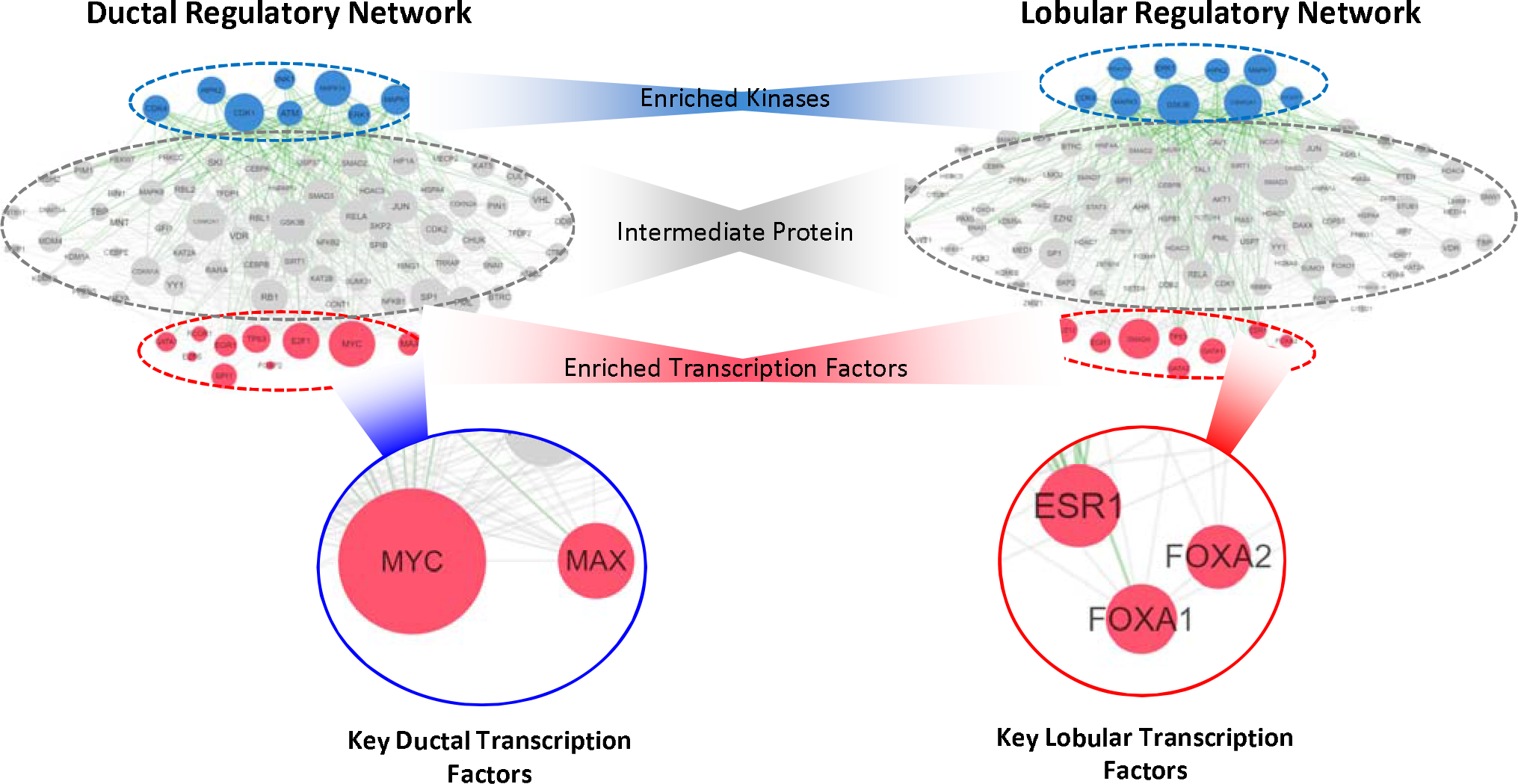
Transcription Factor Enrichment Analysis using Pan-Patient Ductal vs Lobular DEGs. Ductal vs Lobular DEGs were separately input into eXpression2Kinase (X2K) web tool (https://maayanlab.cloud/X2K/) to identify enriched transcription factors and kinases in ductal (left) and lobular (right) tumor regions. The top layer with blue nodes shows kinases, middle layer shows intermediate proteins and bottom layer with red nodes shows transcription factors. An image inset is shown highlighting MYC and ER signaling related transcription factors enriched in ductal and lobular tumor regions, respectively.

**Supplementary Figure S4:**
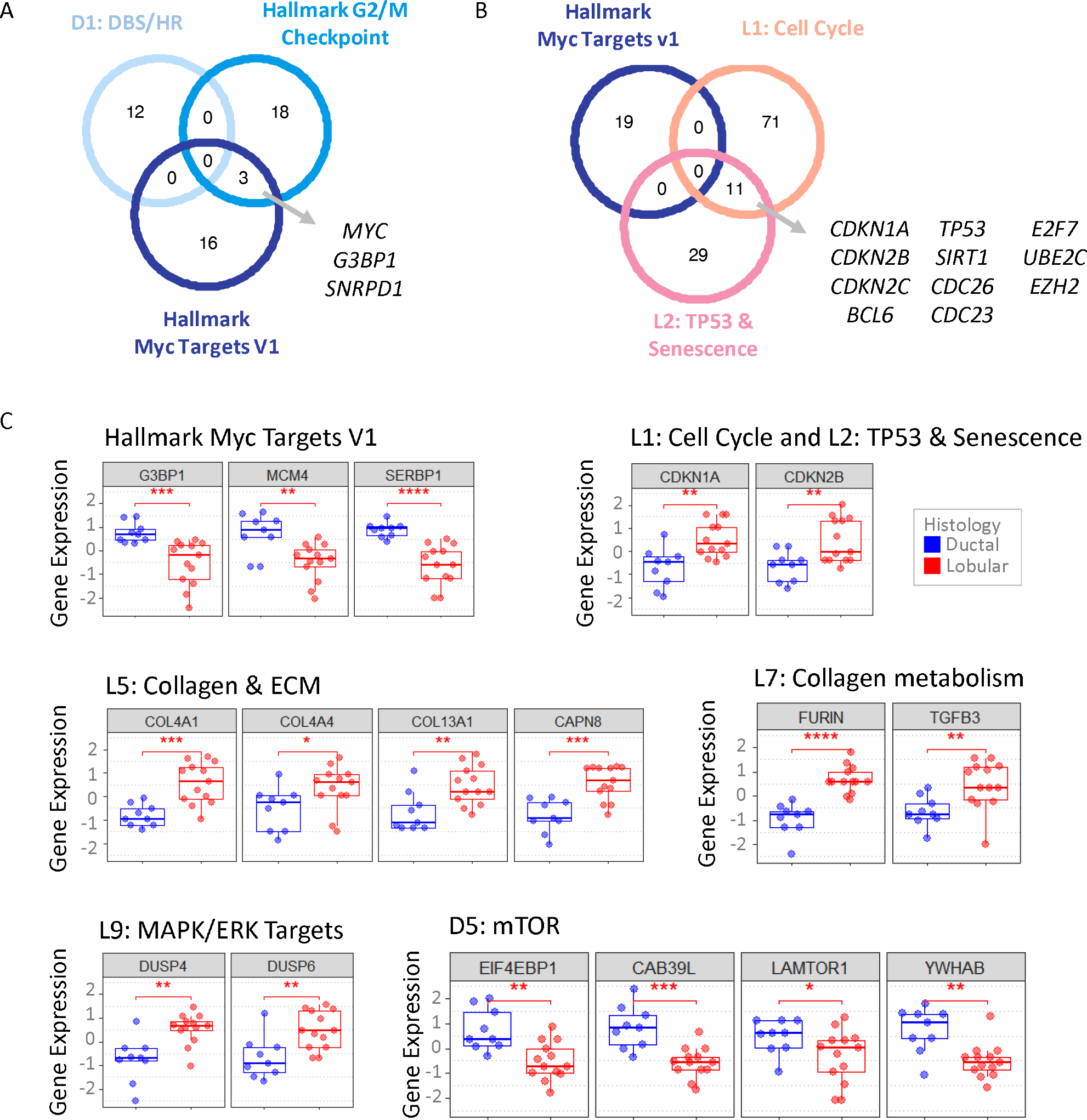
Key Genes Associated with Notable Differentially Enriched Oncogenic and Biological Signatures. (A) Venn diagram showing overlap between D1: Double stranded break/homologous recombination (DBS/HR), Hallmark G2m Checkpoint and Hallmark Myc Targets V1. (B). Venn diagram showing overlap between Hallmark Myc Targets V1, L1: Cell Cycle and L2: TP53 & Senescence. Details of all genes in each signature shown in (A) and (B) are listed in Supplementary Tables S6 and S7. The genes used for comparing biological theme signature clusters (D1: DBS/HR, L1: Cell Cycle and L2: TP53 & Senescence) are defined as union of all individual signatures present in each of these signature clusters. (C) Expression of select genes from Hallmark Myc Targets V1, L1: Cell Cycle, L2: TP53/Senescence, L5: Collagen & ECM, L7: Collagen metabolism, L9: MAPK/ERK Targets, and D5: mTOR in ductal vs. lobular tumor regions. Statistical significance was evaluated using the T test.

**Supplementary Figure S5:**
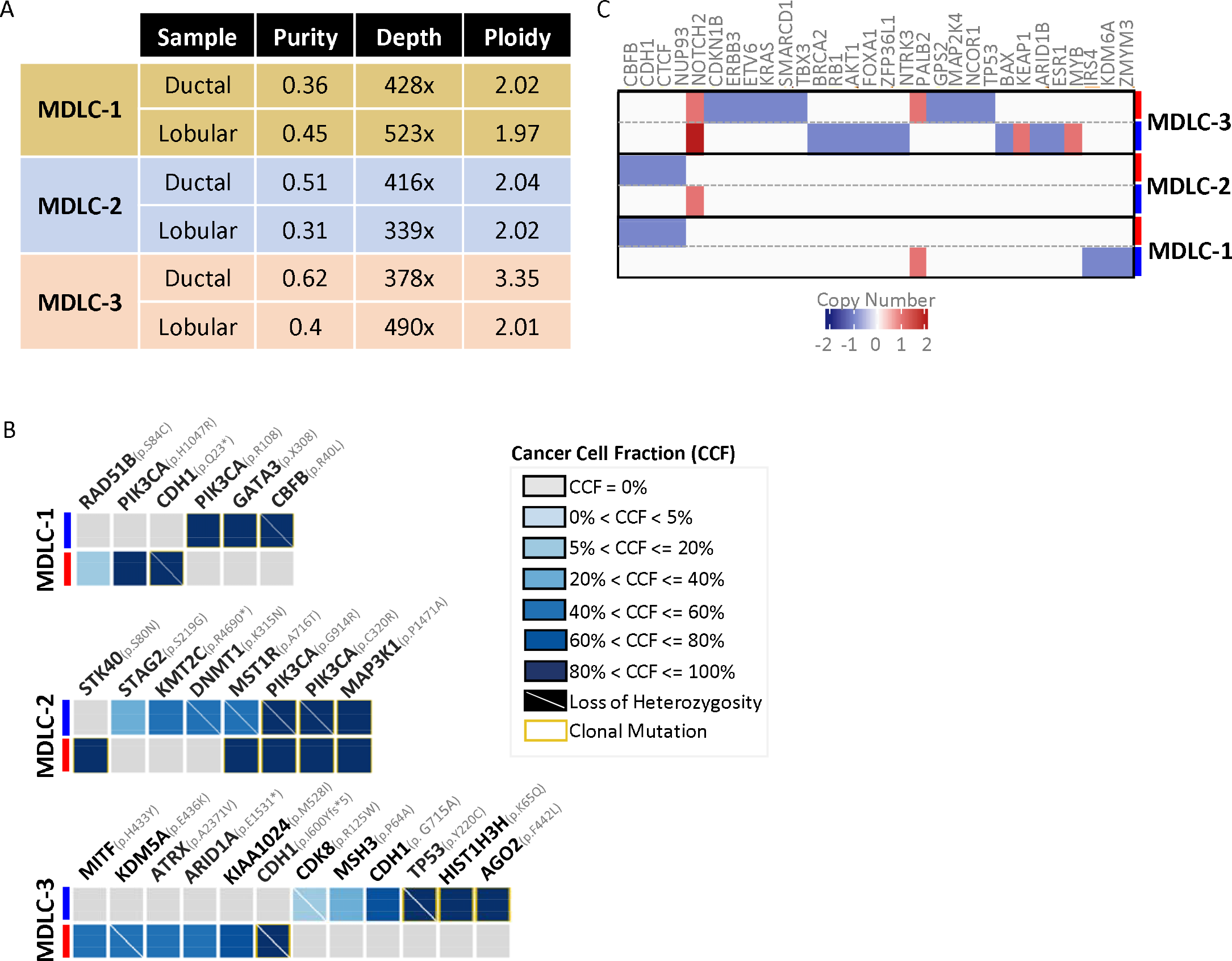
MSK-IMPACT Extended Results. (A) Tumor purity, sequencing coverage and ploidy of ductal and lobular tumor regions across profiled cases. (B) Somatic mutations and their cancer cell fraction (CCF). High CCF indicate clonal mutations while low CCF indicates sub-clonal mutations. (C) Copy number of key breast cancer genes in ductal vs lobular regions across each case.

**Supplementary Figure S6:**
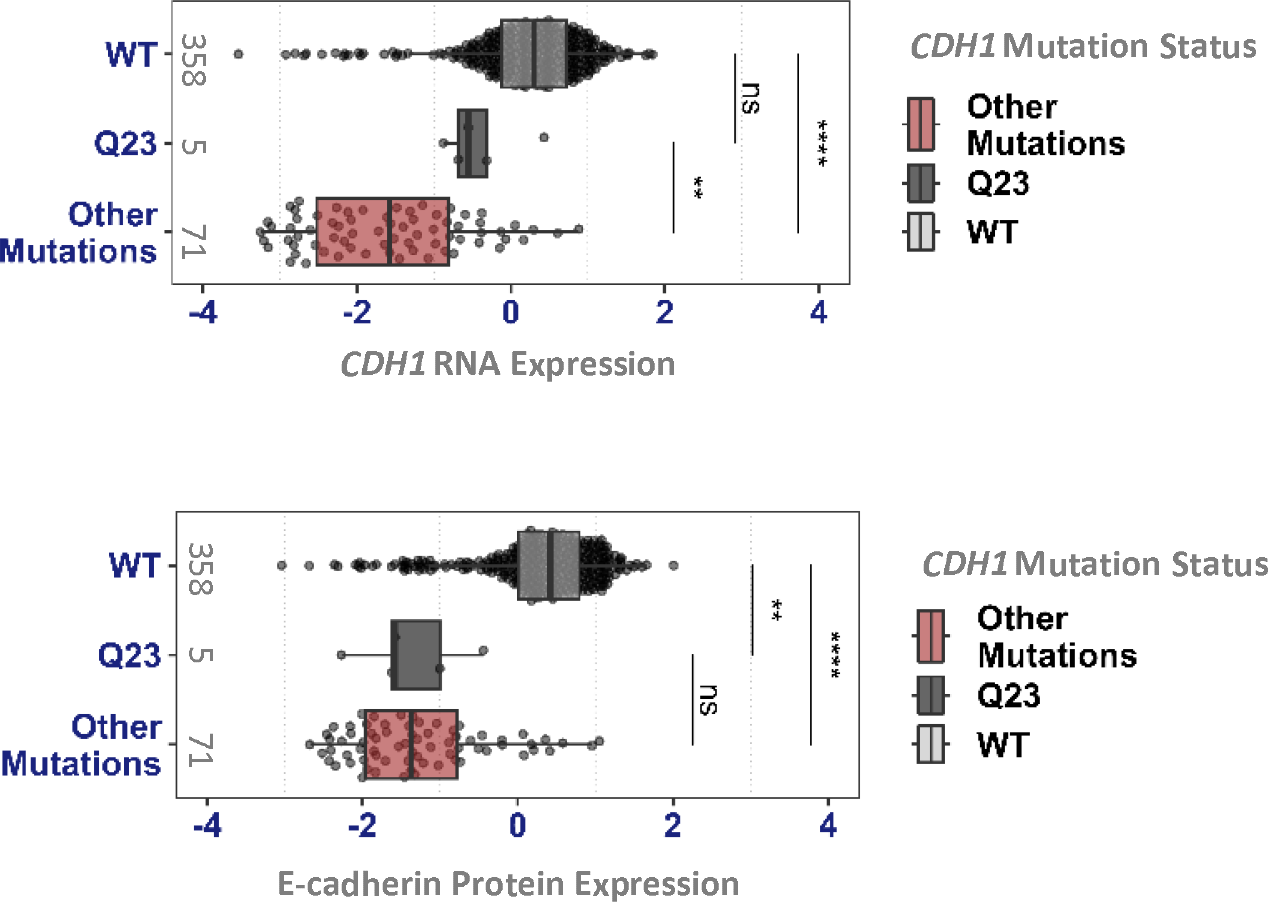
Effect of Q23 mutation on CDH1 RNA and E-cadherin protein expression in ER+ Breast Tumors. Bar plots showing CDH1 RNA (top) and E-cadherin protein (bottom) expression in ER+ patient tumor samples (N=434) with CDH1 Q23* mutations (N=5), all other CDH1 truncating mutations (N=71) and CDH1 WT (N=358) in ER+ breast tumors in TCGA cohort. To avoid confounding effects of CDH1 copy number, we excluded all tumor samples with CDH1 gains, amplification, and deep deletions. Statistical significance is determined using student’s t-test (****: P ≤ 0.0001, **: P ≤ 0.01).

**Supplementary Figure S7:**
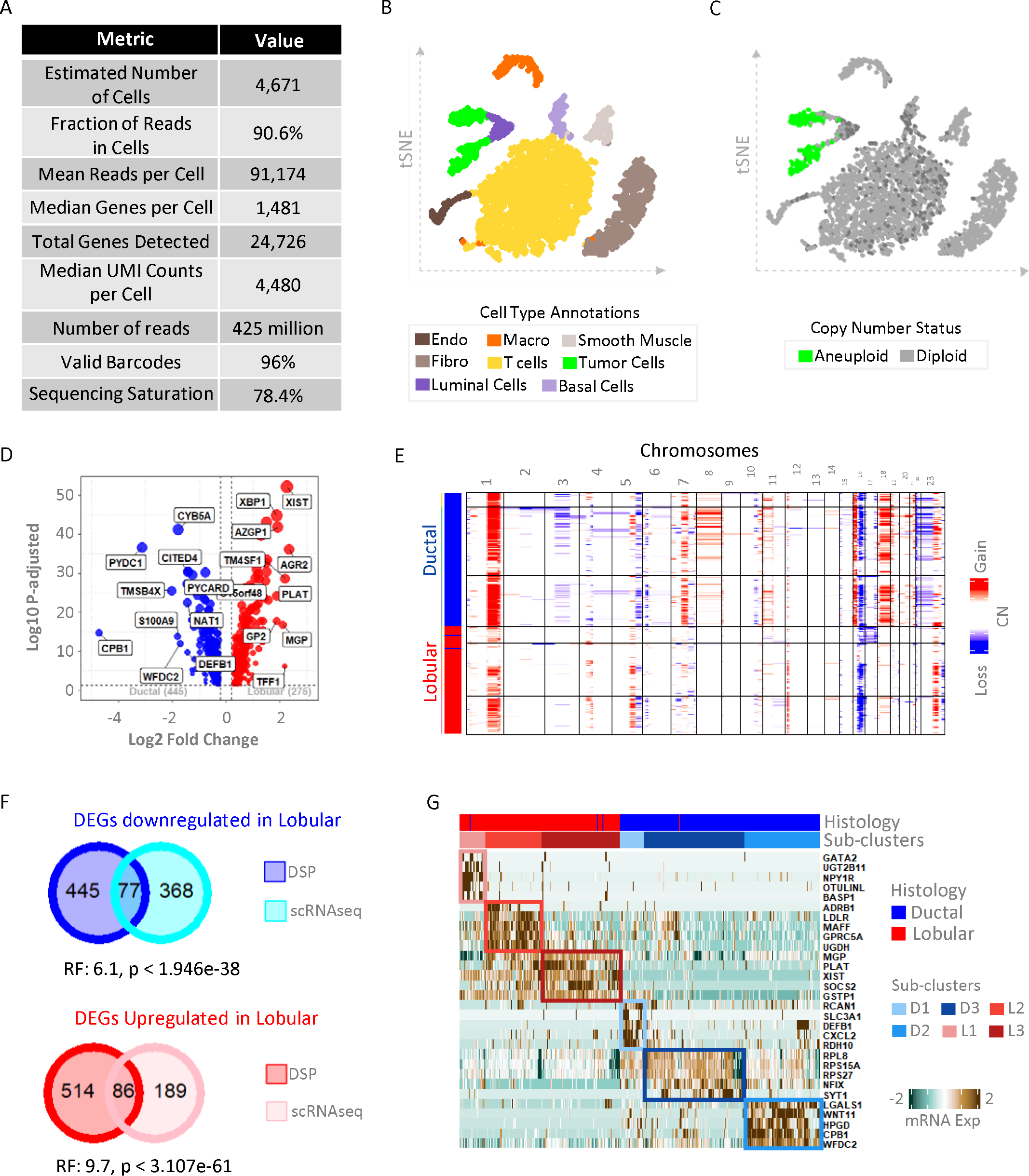
MDLC-1 Ductal and Lobular Tumor scRNAseq Heterogeneity. (A) MDLC-1 single-cell RNAseq (scRNAseq) dataset quality control metrics from cell ranger software. (B) tSNE plot showing identified cell types in the scRNAseq dataset including T-cells, Endothelial cells (Endo), Macrophages (Macro), Fibroblasts (Fibro), Smooth muscle cells, luminal tumor epithelial cells (Tumor cells), luminal epithelial cells (Luminal cells) and basal epithelial cells (Basal cells). (C) CopyKat algorithm predictions for copy number status of scRNAseq dataset to confirm aneuploidy status of tumor cells and diploid status of non-tumor cells. (D) MDLC-1 scRNAseq ductal vs lobular DEG analysis. (E) Heatmap of CopyKat inferred copy number values for ductal and lobular tumor cell populations. (F) Overlap of MDLC-1 ductal vs lobular DEGs between scRNAseq and digital spatial profiling (DSP) datasets. A RF > 1 and p < 0.01 shows significantly strong overlap. Statistical significance of DEG overlap between DEGs from two technologies was computed using overlap_stats tool as described previously in Supplementary Fig. S2. (G) Top five marker genes of each ductal and lobular sub-cluster identified in Fig. 5C.

**Supplementary Figure S8:**
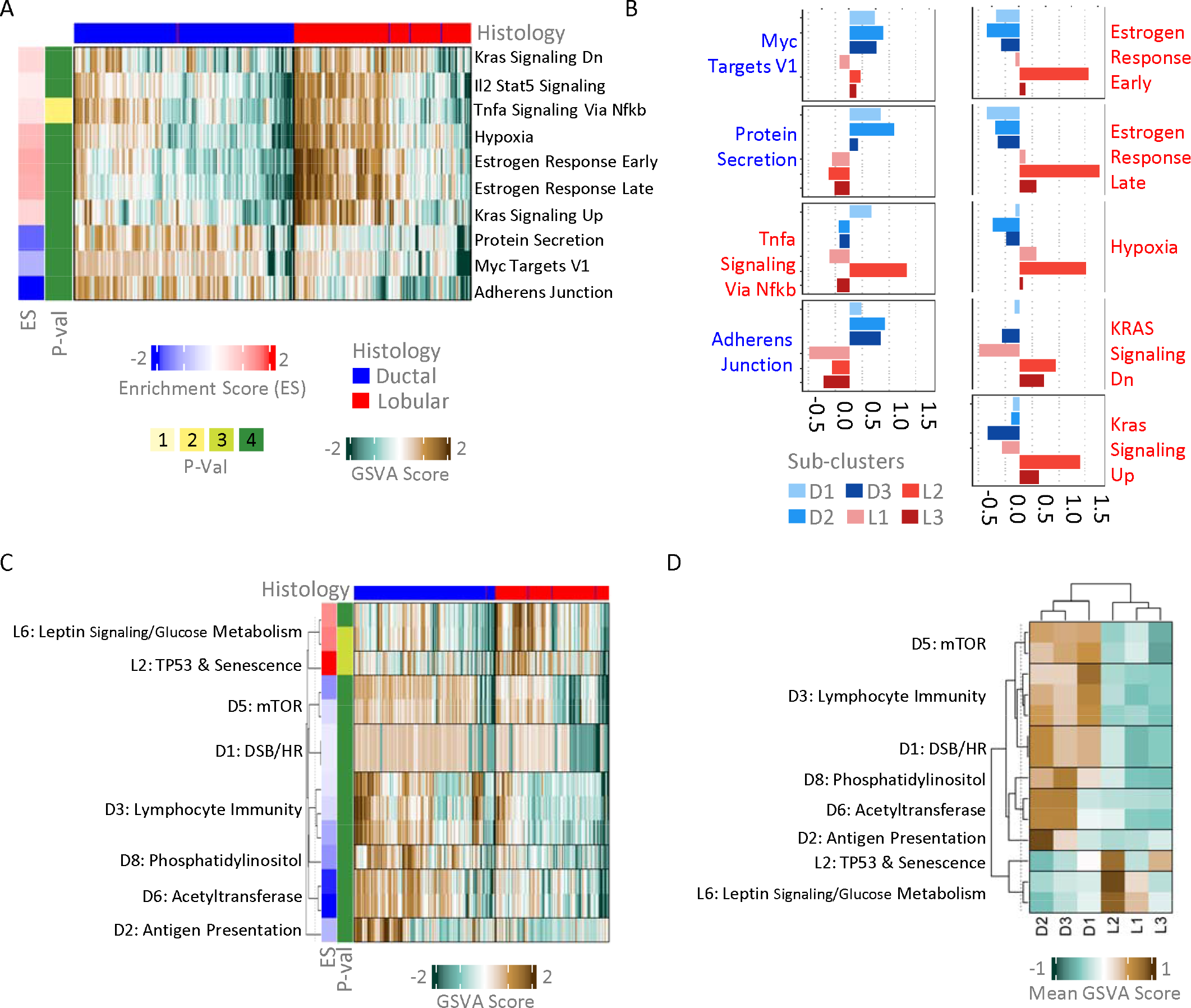
Distinct Biological Signatures in MDLC-1 Ductal and Lobular Tumor scRNAseq. (A) GSVA based hallmark signature enrichment analysis in scRNAseq dataset. Enrichment score (ES) for a pathway shows mean differences of GSVA score for that pathway in lobular – ductal regions and the −log10 of FDR-adjusted P-value (P-val) shows statistical significance of enrichment based on T test. (P-val score definitions: 4 = P ≤ 0.0001, 3 = P ≤ 0.001, 2 = P ≤ 0.01 and 1 = P ≤ 0.05). (B) Barplots showing mean GSVA scores of hallmark signatures in ductal and lobular subclusters. (C) Recurring biology themes in ductal vs lobular cells in scRNAseq dataset. (D) Mean GSVA score for select differentially enriched biology themes across ductal and lobular sub-clusters.

## References

1. Bray, F. et al. Global cancer statistics 2018: GLOBOCAN estimates of incidence and mortality worldwide for 36 cancers in 185 countries. 68, 394–424 (2018).

2. Siegel, R. L., Miller, K. D., Wagle, N. S. & Jemal, A. Cancer statistics, 2023. CA Cancer J Clin 73, 17–48 (2023).

3. Sledge, G. W., Chagpar, A. & Perou, C. Collective wisdom: lobular carcinoma of the breast. Am Soc Clin Oncol Educ B 35, 18–21 (2016).

4. Reed, M. E. M. C., Kutasovic, J. R., Lakhani, S. R. & Simpson, P. T. Invasive lobular carcinoma of the breast: morphology, biomarkers and ’omics. 17, 12 (2015).

5. Ciriello, G. et al. Comprehensive molecular portraits of invasive lobular breast cancer. Cell 163, 506–519 (2015).

6. Reed, M. E. M. C., Kutasovic, J. R., Lakhani, S. R. & Simpson, P. T. Invasive lobular carcinoma of the breast: morphology, biomarkers and ’omics. 17, 12 (2015).

7. Dabbs, D. J., Bhargava, R. & Chivukula, M. Lobular versus ductal breast neoplasms: the diagnostic utility of p120 catenin. Am J Surg Pathol 31, 427–37 (2007).

8. Arpino, G. et al. Infiltrating lobular carcinoma of the breast: tumor characteristics and clinical outcome. 6, (2004).

9. Oesterreich, S. et al. Clinicopathological Features and Outcomes Comparing Patients With Invasive Ductal and Lobular Breast Cancer. J Natl Cancer Inst 114, 1511–1522 (2022).

10. Z, C., et al. Invasive lobular carcinoma of the breast: A special histological type compared with invasive ductal carcinoma. 12, e0182397 (2017).

11. Yeatman, T. J. et al. Tumor biology of infiltrating lobular carcinoma. Implications for management. Ann Surg 222, 549–559 (1995).

12. Helvie, M. A., Paramagul, C., Anderson, K. L. & Adler, D. D. Invasive lobular carcinoma. Imaging features and clinical detection. Invest Radiol 28, 202–207 (1993).

13. Arpino, G. et al. Infiltrating lobular carcinoma of the breast: tumor characteristics and clinical outcome. 6, (2004).

14. Oesterreich, S. et al. Clinicopathological Features and Outcomes Comparing Patients With Invasive Ductal and Lobular Breast Cancer. J Natl Cancer Inst 114, 1511–1522 (2022).

15. Mathew, A. et al. Distinct Pattern of Metastases in Patients with Invasive Lobular Carcinoma of the Breast. Geburtshilfe Frauenheilkd (2017) doi:10.1055/s-0043-109374.

16. Lamovec, J. & Bračkko, M. Metastatic pattern of infiltrating lobular carcinoma of the breast: an autopsy study. 48, 28–33 (1991).

17. Djerroudi, L., Cabel, L., Bidard, F. C. & Vincent-Salomon, A. Invasive Lobular Carcinoma of the Breast: Toward Tailoring Therapy? 114, 1434–1436 (2022).

18. K, V. B., et al. Current and future diagnostic and treatment strategies for patients with invasive lobular breast cancer. Annals of Oncology 33, 769–785 (2022).

19. Luveta, J., Parks, R. M., Heery, D. M., Cheung, K.-L. & Johnston, S. J. Invasive Lobular Breast Cancer as a Distinct Disease: Implications for Therapeutic Strategy. Oncol Ther 8, 1–11 (2020).

20. K, V. B., et al. Current and future diagnostic and treatment strategies for patients with invasive lobular breast cancer. Annals of Oncology 33, 769–785 (2022).

21. Hoon Tan, P., et al. The 2019 WHO classification of tumours of the breast. Histopathology (2020) doi:10.1111/his.14091.

22. Sinn, H. P. & Kreipe, H. A Brief Overview of the WHO Classification of Breast Tumors, 4th Edition, Focusing on Issues and Updates from the 3rd Edition. Breast Care vol. 8 149 (Karger Publishers, 2013).

23. Metzger-Filho, O. et al. The Diagnosis of Mixed Invasive Ductal and Lobular Carcinoma (IDC-L) in Clinical Practice Is Often Associated with Uncertainty Related to Its Prognosis and Response to Systemic Therapies. With the Increasing Recognition of Invasive Lobular Carc. vol. 24 (Oxford University Press (OUP), 2019).

24. Zengel, B. et al. Comparison of the clinicopathological features of invasive ductal, invasive lobular, and mixed (invasive ductal + invasive lobular) carcinoma of the breast. Breast Cancer (2015) doi:10.1007/s12282-013-0489-8.

25. Xiao, Y. et al. Mixed invasive ductal and lobular carcinoma has distinct clinical features and predicts worse prognosis when stratified by estrogen receptor status. Sci Rep (2017) doi:10.1038/s41598-017-10789-x.

26. Duraker, N., Hot, S., Akan, A. & Ozay Nayir, P. A Comparison of the Clinicopathological Features, Metastasis Sites and Survival Outcomes of Invasive Lobular, Invasive Ductal and Mixed Invasive Ductal and Lobular Breast Carcinoma. Eur J Breast Health 16, 22–31 (2020).

27. Nasrazadani, A. et al. Mixed invasive ductal lobular carcinoma is clinically and pathologically more similar to invasive lobular than ductal carcinoma. British Journal of Cancer 2023 1–10 (2023) doi:10.1038/s41416-022-02131-8.

28. Ciriello, G. et al. Comprehensive molecular portraits of invasive lobular breast cancer. Cell 163, 506–519 (2015).

29. Nasrazadani, A. et al. Mixed invasive ductal lobular carcinoma is clinically and pathologically more similar to invasive lobular than ductal carcinoma. British Journal of Cancer 2023 1–10 (2023) doi:10.1038/s41416-022-02131-8.

30. Lohani, K. R. et al. Lobular-Like Features and Outcomes of Mixed Invasive Ductolobular Breast Cancer (MIDLC): Insights from 54,403 Stage I-III MIDLC Patients. Ann Surg Oncol 31, 936–946 (2024).

31. Pérez-Mies, B. et al. The Clonal Relationship Between the Ductal and Lobular Components of Mixed Ductal-Lobular Carcinomas Suggested a Ductal Origin in Most Tumors. American Journal of Surgical Pathology 46, 1545–1553 (2022).

32. Li, C. I., Anderson, B. O., Daling, J. R. & Moe, R. E. Trends in Incidence Rates of Invasive Lobular and Ductal Breast Carcinoma. J Am Med Assoc (2003) doi:10.1001/jama.289.11.1421.

33. Suryadevara, A., Paruchuri, L. P., Banisaeed, N., Dunnington, G. & Rao, K. A. The clinical behavior of mixed ductal/lobular carcinoma of the breast: A clinicopathologic analysis. World J Surg Oncol (2010) doi:10.1186/1477-7819-8-51.

34. Rakha, E. A. et al. The biological and clinical characteristics of breast carcinoma with mixed ductal and lobular morphology. Breast Cancer Res Treat (2009) doi:10.1007/s10549-008-0007-4.

35. Acs, G., Lawton, T. J., Rebbeck, T. R., LiVolsi, V. A. & Zhang, P. J. Differential Expression of E-Cadherin in Lobular and Ductal Neoplasms of the Breast and Its Biologic and Diagnostic Implications. Am J Clin Pathol (2001) doi:10.1309/FDHX-L92R-BATQ-2GE0.

36. McCart Reed, A. E., et al. Mixed ductal-lobular carcinomas: evidence for progression from ductal to lobular morphology. Journal of Pathology (2018) doi:10.1002/path.5040.

37. Christgen, M. et al. Inter-observer agreement for the histological diagnosis of invasive lobular breast carcinoma. J Pathol Clin Res 8, 191–205 (2022).

38. Metzger Filho, O., et al. Mixed Invasive Ductal and Lobular Carcinoma of the Breast: Prognosis and the Importance of Histologic Grade. Oncologist 24, (2019).

39. Zhao, H. The prognosis of invasive ductal carcinoma, lobular carcinoma and mixed ductal and lobular carcinoma according to molecular subtypes of the breast. Breast Cancer 28, 187–195 (2021).

40. Arps, D. P., Healy, P., Zhao, L., Kleer, C. G. & Pang, J. C. Invasive ductal carcinoma with lobular features: a comparison study to invasive ductal and invasive lobular carcinomas of the breast. Breast Cancer Res Treat 138, 719 (2013).

41. Zels, G. et al. Metastases of primary mixed no-special type and lobular breast cancer display an exclusive lobular histology. Breast 75, 103732 (2024).

42. M, C., et al. Lobular breast cancer: Histomorphology and different concepts of a special spectrum of tumors. 13, 3695 (2021).

43. Christgen, M., Kandt, L. D., Antonopoulos, W. & al., et. Inter-observer agreement for the histological diagnosis of invasive lobular breast carcinoma. J Pathol Clin Res 8, 191–205 (2022).

44. Tang, X. et al. Mixed ductal–lobular carcinoma: an analysis of CDH1 DNA copy number variation and mutation. Breast Cancer 28, 1318–1327 (2021).

45. McCart Reed, A. E., et al. Mixed ductal-lobular carcinomas: evidence for progression from ductal to lobular morphology. Journal of Pathology (2018) doi:10.1002/path.5040.

46. Yu, J. et al. Clinicopathologic and genomic features of lobular like invasive mammary carcinoma: is it a distinct entity? npj Breast Cancer 2023 9:1 9, 1–13 (2023).

47. Ciriello, G. et al. Comprehensive Molecular Portraits of Invasive Lobular Breast Cancer. Cell 163, 506–519 (2015).

48. Suryadevara, A., Paruchuri, L. P., Banisaeed, N., Dunnington, G. & Rao, K. A. The clinical behavior of mixed ductal/lobular carcinoma of the breast: A clinicopathologic analysis. World J Surg Oncol (2010) doi:10.1186/1477-7819-8-51.

49. Pérez-Mies, B. et al. The Clonal Relationship Between the Ductal and Lobular Components of Mixed Ductal-Lobular Carcinomas Suggested a Ductal Origin in Most Tumors. American Journal of Surgical Pathology 46, 1545–1553 (2022).

50. Liberzon, A. et al. The Molecular Signatures Database Hallmark Gene Set Collection. Cell Syst 1, 417–425 (2015).

51. Carbon, S. et al. The Gene Ontology resource: enriching a GOld mine. Nucleic Acids Res 49, D325–D334 (2021).

52. Gillespie, M. et al. The reactome pathway knowledgebase 2022. Nucleic Acids Res 50, D687–D692 (2022).

53. Kanehisa, M., Furumichi, M., Sato, Y., Kawashima, M. & Ishiguro-Watanabe, M. KEGG for taxonomy-based analysis of pathways and genomes. Nucleic Acids Res 51, D587– D592 (2023).

54. Cleton-Jansen, A. M. E-cadherin and loss of heterozygosity at chromosome 16 in breast carcinogenesis: different genetic pathways in ductal and lobular breast cancer? Breast Cancer Res 4, 5–8 (2002).

55. André, F. et al. Alpelisib for PIK3CA -Mutated, Hormone Receptor–Positive Advanced Breast Cancer. New England Journal of Medicine 380, 1929–1940 (2019).

56. Theodorou, V., Stark, R., Menon, S. & Carroll, J. S. GATA3 acts upstream of FOXA1 in mediating ESR1 binding by shaping enhancer accessibility. Genome Res 23, 12 (2013).

57. Malik, N. et al. The transcription factor CBFB suppresses breast cancer through orchestrating translation and transcription. Nature Communications 2019 10:1 10, 1–15 (2019).

58. Comprehensive molecular portraits of human breast tumours. Nature 490, (2012).

59. Gala, K. et al. KMT2C mediates the estrogen dependence of breast cancer through regulation of ERα enhancer function. Oncogene 2018 37:34 37, 4692–4710 (2018).

60. Glaser, A. P. et al. APOBEC-mediated mutagenesis in urothelial carcinoma is associated with improved survival, mutations in DNA damage response genes, and immune response. Oncotarget 9, 4537–4548 (2017).

61. Shen, R. & Seshan, V. E. FACETS: allele-specific copy number and clonal heterogeneity analysis tool for high-throughput DNA sequencing. Nucleic Acids Res 44, (2016).

62. Glaser, A. P. et al. APOBEC-mediated mutagenesis in urothelial carcinoma is associated with improved survival, mutations in DNA damage response genes, and immune response. Oncotarget 9, 4537–4548 (2017).

63. Klonowski, J., Liang, Q., Coban-Akdemir, Z., Lo, C. & Kostka, D. aenmd: annotating escape from nonsense-mediated decay for transcripts with protein-truncating variants. Bioinformatics 39, (2023).

64. Khajavi, M., Inoue, K. & Lupski, J. R. Nonsense-mediated mRNA decay modulates clinical outcome of genetic disease. European Journal of Human Genetics 2006 14:10 14, 1074–1081 (2006).

65. Litchfield, K. et al. Escape from nonsense-mediated decay associates with anti-tumor immunogenicity. Nature Communications 2020 11:1 11, 1–11 (2020).

66. Moffitt, J. R., Lundberg, E. & Heyn, H. The emerging landscape of spatial profiling technologies. Nature Reviews Genetics 2022 23:12 23, 741–759 (2022).

67. van Amerongen, R. Behind the Scenes of the Human Breast Cell Atlas Project. J Mammary Gland Biol Neoplasia 26, 67 (2021).

68. Nguyen, Q. H. et al. Profiling human breast epithelial cells using single cell RNA sequencing identifies cell diversity. Nat Commun (2018) doi:10.1038/s41467-018-04334-1.

69. Gray, G. K. et al. A human breast atlas integrating single-cell proteomics and transcriptomics. Dev Cell 57, 1400–1420.e7 (2022).

70. Wu, S. Z. et al. A single-cell and spatially resolved atlas of human breast cancers. Nature Genetics 2021 53:9 53, 1334–1347 (2021).

71. Sun, H. et al. The relevance between hypoxia-dependent spatial transcriptomics and the prognosis and efficacy of immunotherapy in claudin-low breast cancer. Front Immunol 13, 7828 (2023).

72. Bassiouni, R. et al. Spatial Transcriptomic Analysis of a Diverse Patient Cohort Reveals a Conserved Architecture in Triple-Negative Breast Cancer. Cancer Res 83, 34–48 (2023).

73. Johnson, B. E. et al. An omic and multidimensional spatial atlas from serial biopsies of an evolving metastatic breast cancer. Cell Rep Med 3, 100525 (2022).

74. Kulasinghe, A. et al. Spatial Profiling Identifies Prognostic Features of Response to Adjuvant Therapy in Triple Negative Breast Cancer (TNBC). Front Oncol 11, 1 (2021).

75. Andersson, A. et al. Spatial deconvolution of HER2-positive breast cancer delineates tumor-associated cell type interactions. Nature Communications 2021 12:1 12, 1–14 (2021).

76. Badve, S. et al. Basal-like and triple-negative breast cancers: a critical review with an emphasis on the implications for pathologists and oncologists. Modern Pathology 24, 157–167 (2011).

77. Perou, C. M. Molecular Stratification of Triple Negative Breast Cancers. Oncologist 16, 61–70 (2011).

78. Michaut, M. et al. Integration of genomic, transcriptomic and proteomic data identifies two biologically distinct subtypes of invasive lobular breast cancer. Sci Rep (2016) doi:10.1038/srep18517.

79. Djerroudi, L., Cabel, L., Bidard, F. C. & Vincent-Salomon, A. Invasive Lobular Carcinoma of the Breast: Toward Tailoring Therapy? 114, 1434–1436 (2022).

80. Gulhan, D. C., Lee, J. J. K., Melloni, G. E. M., Cortés-Ciriano, I. & Park, P. J. Detecting the mutational signature of homologous recombination deficiency in clinical samples. Nat Genet 51, 912–919 (2019).

81. Alexandrov, L. B. et al. Signatures of mutational processes in human cancer. Nature 500, 415–421 (2013).

82. Alexandrov, L. B. et al. The repertoire of mutational signatures in human cancer. Nature 578, 94–101 (2020).

83. Vahdatinia, M. et al. KIT genetic alterations in breast cancer. J Clin Pathol (2022) doi:10.1136/JCP-2022-208611.

84. Geyer, F. C. et al. Genetic analysis of a morphologically heterogeneous ovarian endometrioid carcinoma. Histopathology 71, 480 (2017).

85. Azim, H. A., Nguyen, B., Brohée, S., Zoppoli, G. & Sotiriou, C. Genomic aberrations in young and elderly breast cancer patients. BMC Med 13, 1–7 (2015).

86. Afzaljavan, F., Sadr, A. S., Savas, S. & Pasdar, A. GATA3 somatic mutations are associated with clinicopathological features and expression profile in TCGA breast cancer patients. Scientific Reports 2021 11:1 11, 1–13 (2021).

87. BeLow, M. & Osipo, C. Notch Signaling in Breast Cancer: A Role in Drug Resistance. Cells 2020, Vol. 9, Page 2204 9, 2204 (2020).

88. Hopkins, J. L., Lan, L. & Zou, L. DNA repair defects in cancer and therapeutic opportunities. Genes Dev 36, 278 (2022).

89. Voutsadakis, I. A. & Stravodimou, A. Homologous Recombination Defects and Mutations in DNA Damage Response (DDR) Genes Besides BRCA1 and BRCA2 as Breast Cancer Biomarkers for PARP Inhibitors and Other DDR Targeting Therapies. Anticancer Res 43, 967–981 (2023).

90. McCart Reed, A. E., et al. An epithelial to mesenchymal transition programme does not usually drive the phenotype of invasive lobular carcinomas. Journal of Pathology 238, 489–494 (2016).

91. Herschkowitz, J. I. et al. Identification of conserved gene expression features between murine mammary carcinoma models and human breast tumors. Genome Biol 8, 1–17 (2007).

92. Pan, C. et al. Research progress of Claudin-low breast cancer. Front Oncol 13, 1226118 (2023).

93. Dias, K. et al. Claudin-Low Breast Cancer; Clinical & Pathological Characteristics. PLoS One 12, (2017).

94. McCart Reed, A. E., et al. Mixed ductal-lobular carcinomas: evidence for progression from ductal to lobular morphology. Journal of Pathology (2018) doi:10.1002/path.5040.

95. Kutasovic, J. R., McCart Reed, A. E., Sokolova, A., Lakhani, S. R. & Simpson, P. T. Morphologic and genomic heterogeneity in the evolution and progression of breast cancer. Cancers vol. 12 Preprint at 10.3390/cancers12040848 (2020).

96. Ciriello, G. et al. Comprehensive Molecular Portraits of Invasive Lobular Breast Cancer. Cell (2015) doi:10.1016/j.cell.2015.09.033.

97. Bajrami, I. et al. E-Cadherin/ROS1 Inhibitor Synthetic Lethality in Breast Cancer. Cancer Discov 8, 498–515 (2018).

98. Jiang, Y., Yang, M., Wang, S., Li, X. & Sun, Y. Emerging role of deep learning-based artificial intelligence in tumor pathology. Cancer Commun 40, 154–166 (2020).

99. Couture, H. D. et al. Image analysis with deep learning to predict breast cancer grade, ER status, histologic subtype, and intrinsic subtype. npj Breast Cancer 2018 4:1 4, 1–8 (2018).

100. Chan, R. C. K. et al. Artificial intelligence in breast cancer histopathology. Histopathology 82, 198–210 (2023).

101. Gerdes, M. J. et al. Highly multiplexed single-cell analysis of formalin-fixed, paraffin-embedded cancer tissue. Proc Natl Acad Sci U S A 110, 11982–11987 (2013).

102. Pachitariu, M. & Stringer, C. Cellpose 2.0: how to train your own model. Nat Methods 19, 1634–1641 (2022).

103. Van Der Walt, S. et al. scikit-image: image processing in Python. PeerJ 2, (2014).

