## Supplementary Methods for "Spatial molecular profiling of mixed invasive ductal-lobular breast cancers reveals heterogeneity in intrinsic molecular subtypes, oncogenic signatures, and mutations"

#### Digital Spatial Profiling (DSP)

DSP was performed by NanoString GeoMX as described previously^1^. Briefly, five-µm-thick FFPE tumor sections were mounted on single positively charged slide. Slides underwent deparaffinization, rehydration, heat-induced epitope retrieval, and enzymatic digestion. Oligonucleotide probe mix from whole-transcriptome atlas (WTA) panel were hybridized and incubated overnight. The slide was blocked and incubated with fluorescent antibodies against morphology markers and DNA dye (Syto83 dye; Invitrogen Cat # S11364) for 1 hour. Morphology markers included E-cadherin (BD Pharmingen Cat# 560062), Pan-cytokeratin (Novus Cat# NBP2-33200AF488) and Vimentin (Santa Cruz Cat# sc-373717 AF594). Finally, the slide was loaded into the GeoMx platform and scanned for an immunofluorescent signal. Together with the aid of H&E and morphology marker stains, our panel of pathologist selected 26 rectangle ROIs (12 lobular ROIs, 11 ductal ROIs and 3 mixed ROIs) with an area of 518,100 μm^2^. To profile ductal and lobular cells separately the following segmentation approach was used; 1) ductal cells a E-cadherin +ive & Pan- cytokeratin +ive, and 2) lobular cells a E-cadherin -ive & Pan- cytokeratin +ive. Above segmentation resulted in total of 29 areas (14 ductal and 15 lobular). UV light was directed at each segmented region to release the oligonucleotides that were collected and then sequenced using Illumina NextSeq 2000. The resulting FASTQ files were processed using GeoMX NGS pipeline 2.3.4 which provided the count data for all target probe across profiled regions. All 29 profiled regions had high sequencing saturation (> 94%) and low signal-to-noise ratio (Q3 count/NegProbe count ≥ 1) (Supplementary Fig. S1A-C; Supplementary Table S2,S3). Further downstream analysis is described below.

##### E-cadherin Quantification

To quantify E-cadherin fluorescence, the segmented regions were analyzed using ImageJ^2^. First, images were converted to 8-bit format. The "Intermodes" color threshold was used to remove the background and retain the fluorescence signal. E-cadherin intensity was measured as the percentage area with a fluorescence signal. To obtain the normalized E-cadherin intensity for each segmented region, the E-cadherin intensity was multiplied by the nuclei count.

##### Quality Control

Three areas of profiled regions were excluded due to low surface area (< 5000 µm^2^), low nuclei count (< 80) and low read count (< 0.3 million). The final DSP dataset is comprised of 13 ductal and 13 lobular regions. Q3 normalized counts were used for all downstream analyses. All downstream analyses were performed in R^3^ unless specified otherwise. Additional information on properties of profiled regions including surface area, nuclei count, and sequencing metrics is provided in Supplementary Table S2. Q3 normalized counts are provided in Supplementary Table S3.

##### Data Exploration & Transcriptomic Clusters

Transcriptomic similarity across profiled regions was assessed by tSNE^4,5^ and hierarchical clustering^6^ using top 10% most variable genes. To identify robust transcriptomic classes, consensus clustering approach was employed using ConsensusClusterPlus v1.60.0^7^. PAM50 intrinsic molecular subtypes were computed by genefu v2.28.0^8^ (Supplementary Table S4).

##### Differential Analysis

Differential expression analysis (DEA) between ER+ ductal (N=9) vs lobular (N=13) regions was performed using Limma Voom v3.52.2^9,10^ with the following design: ~Histology [i.e., ductal or lobular] + PatientID. Note: Genes with +ive log fold change (LFC) show an up-regulation in the ductal vs lobular regions and vice versa. Significant differentially expressed genes (DEG) were defined as those with absolute LFC > 0.25 and FDR-adjusted P value of ≤ 0.3. Patient-wise DEA between ductal and lobular regions was also done. All patient-wise DEA results were compared to each other and ER+ DEA (Supplementary Fig. S2). All DEA results are listed in Supplementary Table S5. To assess similarity of mDLC ductal and lobular regions with its pure counterparts (i.e., IBC-NST and ILC), we compared mDLC ductal vs. lobular DEG with those from TCGA IBC-NST vs ILC DEA analysis. Comparison with TCGA: TCGA IBC-NST vs ILC DEA analysis was computed using the RNAseq data from The Cancer Genome Atlas (TCGA)^11^ (GEO: GSE62944^12^). Importantly, the tumor purity estimates were used as covariates in the design for the TCGA IBC-NST vs ILC DEA analysis to adjust for tumor purity differences between IBC-NST vs ILC tumor samples as described previously^13^.

##### Pathway Analyses

Differential enrichment of Hallmark signatures: Each RNAseq sample was scored for Hallmark signatures (including Hallmark^14^ and KEGG^15^ Adherens Junction genesets in Molecular Signature Database (MSigDB)^16^. Genesets were downloaded using hypeR v1.12.0^17^. Each geneset was filtered to only include genes overlapping with the DEG from ductal vs lobular differential analyses. Geneset variation analysis (GSVA) v1.44.3^18^ was used for scoring each signature (i.e., filtered geneset). Enrichment Score (ES) for each signature was defined as the difference between mean GSVA score of that signature in ductal and lobular regions (in DSP dataset) or clusters (in scRNAseq data). Enrichment p-value was computed by comparing GSVA scores between these groups using student t-test. Signatures with p-value ≤ 0.05 were considered significant (Supplementary Table S6). Comparison with TCGA: Hallmark signature analysis was performed of TCGA ER+ IBC-NST vs ILC tumor samples was also performed as described above. Briefly, hallmark genesets from MSigDB were filtered to only include genes overlapping with the DEG from TCGA ER+ IBC-NST vs ILC differential analyses using DeSeq2. GSVA was used for scoring each signature (i.e., filtered geneset) in TCGA ER+ IBC-NST and ILC samples. ES and Enrichment p-value were computed as defined previously.

Biological themes: For identifying recurring biological themes in DSP dataset, the differential enrichment analysis described above was expanded to include all GO^19,20^, REACTOME^21^, and KEGG genesets. Signatures and their ES and p-value were computed as aforementioned. Signatures with p-value ≤ 0.05 were shortlisted. Jaccard index (JI) similarity matrix for all significant signatures was computed. All signatures with strong similarity (JI ≥ 0.5) with at least two other signatures were grouped together and manually annotated based on the biological terms used in their original names. For a simplified summary of key recurrent biological themes that were also clinically meaningful, we selected signatures containing following biological terms: Collagen, ECM, Extracellular Matrix, Ncam1, Immunoglobulin, Mediated Immunity, Leptin, Glucose Metabolic, Gluconeogenesis, Mitotic, Cell Cycle, Senescence, Proteoglycan, Tp53, Proteosome, Mhc Class I, Antigen Processing, Oxidative Phosphorylation, Electron Transport Chain, Respiration, Sumoylation, Acetyltransferase, Oxidative Stress, Double Strand, DNA Repair, Mtor, Phosphatidylinositol, Protein Maturation, Unfolded Protein Response and Mapk Targets. Robustness of each significant signature was assessed against background permutations. Signatures that were significant in background permutations were considered not robust and removed from further analysis. Background permutations were computed as following; GSVA was reperformed on all genesets after filtering to only include genes overlapping with a ‘background’ gene list. ‘Background’ gene list was generated by random sampling of non-DEG gene space (i.e., all genes excluding the ones in DEG list from ductal vs lobular differential analysis of DSP dataset). The size of each ‘background’ gene list was same as the DEG list. The above approach was repeated 5000 times (i.e., GSVA was performed each time on all genesets after they were filtered using a sampled ‘background’ gene list. Any signature that was significant between ductal and lobular groups comparisons in more than 5% of all ‘background’ gene list permutations was excluded. The key recurring biological themes and associated signatures are listed in Supplementary Table S7. Transcription factor and kinase enrichment analysis: DEG list were divided into lobular genes (up-regulated in ductal vs lobular regions) and ductal genes (down-regulated in ductal vs lobular regions). Each gene set was individually analyzed in expressionX2Kinases webtools (<https://maayanlab.cloud/X2K/>).

#### MSK-Impact Mutation Profiling

Eight-μm-thick sections from representative FFPE tumor and matched normal tissue sections from MDLC-3, MDLC-2 & MDLC-1 were stained with nuclear fast red and microdissected using a sterile needle under a stereomicroscope (Olympus SZ61) to enrich tumor content, as previously described^22,23^. Histologically distinct tumor components were microdissected separately. Genomic DNA of each component and of matched normal tissue was extracted using the DNeasy Blood and Tissue Kit (Qiagen) according to manufacturers’ instructions. Tumor and normal DNA from each case were subjected to massively parallel sequencing using the Food and Drug Administration (FDA) approved Memorial Sloan Kettering Integrated Mutation Profiling of Actionable Cancer Targets (MSK-IMPACT) multigene panel targeting all coding regions of 515 cancer-related genes as previously described^24–26^. Somatic alterations were identified as previously described^27,28^ (Supplementary Table S8). The median depth of coverage of tumor and normal samples was 416x (range 287–563) and 210x (range 175-217), respectively (Supplementary Fig. S5A). In brief, reads were aligned to the reference human genome GRCh37 using the BWA v0.7.15^29^. Local realignment, duplicate removal and base quality recalibration were performed using the GATK. V3.1.1^30^. Further downstream analysis is described below.

##### Genomic Alteration Analysis

Somatic single nucleotide variants (SNVs.) were detected by MuTect (v1.0)^31^, insertions and deletions (indels) by Strelka^32^, Varscan2^33^, Scalpel^34^ and Lancet^35^. All mutations were manually inspected using the Integrative Genomics Viewer (IGV), and mutations targeting hotspot loci were assigned according to Chang *et al.*^36^. Somatic mutations identified in one histologic component from a given tumor were subsequently interrogated in the matched other histologic component by manual inspection of BAM files using mpileup files (SAMtools mpileup; v1.2 htslib 1.2.1)^37^. Copy number alterations (CNAs) and LOH were identified using FACETS^38^. The cancer cell fraction (CCF) and clonal probability of each mutation was determined using ABSOLUTE^39^. Mutational signatures were inferred using Signature Multivariate Analysis (SigMA), an algorithm for robustly inferring mutational signatures from multigene panel-based sequencing of clinical specimens^40–44^. The shared origin in the disease evolution trees was defined based on presence of overlapping mutations between ductal and lobular regions for each case.

#### *CDH1* promoter methylation assessment by digital droplet PCR

Following PicoGreen quantification, 0.2-9 ng bisulfite treated genomic DNA was combined with locus-specific primers targeting the two *CDH1* promoter CpG islands, FAM- and HEX-labeled probes (Supplementary Table S9), the restriction enzyme HaeIII, and digital PCR Supermix for probes (no dUTP). CpG Methylated DNA (ThermoFisher, Waltham, MA) and Universal Unmethylated DNA (Millipore, Burlington, MA) was used as positive and negative controls, respectively. All reactions were performed on a QX200 ddPCR system (Bio-Rad, Hercules, CA), and each sample was evaluated in two technical duplicates. Reactions were partitioned into ~41K droplets per well using the QX200 droplet generator. Emulsified PCRs were run on a 96-well thermal cycler using the following cycling conditions: 95°C 10’; 50 cycles of 94°C 1’ and 54°C 2’; 98°C 10’. Plates were read and analyzed using the QuantaSoft software (Bio-Rad, Hercules, CA) to assess the number of droplets positive for *CDH1* promoter methylated, unmethylated, both, or neither. Methylation Frequency (MF) was inferred as MF = 100 * Methylated /(Methylated + Unmethylated). Methylation of the *CDH1* promoter was defined as higher than 35 methylated droplets.

#### Single cell RNA sequencing

##### Single Cell Dissociation

Fresh tumor sample underwent single cell dissociation following 10x genomics tumor dissociation protocol (Version: RevB). Briefly, 0.4g tumor tissue was cleaned and minced in serum free DMEM media. Cell dissociation was performed using a combination of both mechanical force and enzymatic digestion using Miltenyi tumor dissociation system (Miltenyi: 130-093-235, 130-095-929) as per manufacturer’s protocol. Next the cell suspension with remaining tissue was filtered through 70 µM strainers (Fisher: 08-771-2). Red blood cells lysed was performed using RBC lysis buffer (Qiagen: 158904). Cellular viability was measured by trypan blue staining. Dead cell removal was performed using dead cell removal kit (Miltenyi: 130-090-101) to acquire around 8000 cells with viability of > 65%.

##### Library Preparation & Sequencing

Freshly dissociated cells were used for scRNAseq library preparation using the 10X genomics V3 chemistry 3’ end protocol. The cells were washed with PBS+0.04% BSA once and then resuspended into 1000-1200 cells/µl to generate Gel Beads in Emulsion (GEMs). Reverse transcription was performed, and the cDNA released from GEMS was purified by DynaBeads, PCR amplified before library preparation. cDNA fragment size quality control was performed using Agilent TapeStation. scRNAseq libraries were sequenced at UPMC Genome Center with NovaSeq 6000 with a depth of 100K reads per cell with setting read1-28bp, read2-91bp, and i7 index-8bp.

##### scRNAseq Analysis

Raw BCL files from Novaseq6000 were converted to FASTQ files with Cell Ranger (Version: 3.2.0) mkfastq function. Cell Ranger count function was then applied for read alignment, barcode and UMI counting using GRCh38 as reference. In summary, we got high quality data: 4,671 cells with 90% reads in cells and mean of 91K reads per cell (Supplementary Fig. S7A). All downstream analyses were performed in R. Seurat v4.1.1^45^ was used for data normalization, dimension reduction and clustering. Briefly, cells with less than 200 genes and genes found in less than 3 ells were removed. Gene counts were Log normalized and scaled. tSNE was used for dimensionality reduction and visualization. Various cell types including were identified using previously published canonical markers^46,47^. Briefly, the following cell types were identified using their respective markers: luminal epithelial cells {*KRT7, KRT8, KR18, KRT19*}, basal epithelial cells {*KRT15, VIM*}, tumor epithelial cells {*ESR1, CDH1, EPCAM, GATA3*}, Endothelial cells {*CDH5, VWF*}, Fibroblasts {*HTRA1, FAP, FBN1*}, Smooth muscle cells {*MYLK, MYL9, ACTA2*}, Macrophages {*CSF1R, CD68, CD163, CD14, PTPRC*} and T-cells {*CD3D, PTPRC*}. To confirm identity of tumor epithelial cells, copy number analysis was performed using CopyKat v1.0.8^48^ and aneuploid tumor epithelial cells were annotated. Tumor epithelial cells were re-clustered using the lowest clustering resolution (0.1) (Fig. 5A). Ductal and lobular scRNAseq clusters were identified by scoring for DSP derived ductal and lobular region signatures (listed in Supplementary Table S5 – comparison 4 column 5) using Seurat’s “AddModuleScore” function. DEGs were identified between ductal and lobular scRNAseq clusters using Seurat’s “FindMarker” function (Supplementary Fig. S7B, Supplementary Table S10). Significant DEGs were defined as absolute LFC > 0.25 and p-adjusted ≤ 0.05. Overlap of scRNAseq DEGs with DSP DEGs is shown in Supplementary Fig. S7C. Significance of the overlap was tested using the overlap_stats tool (http://nemates.org/MA/progs/overlap_stats.html). Hallmark signature analysis, as described above in DSP methods, was performed on scRNAseq ductal vs. lobular cluster DEGs (Supplementary Fig. S7D) and was extended to the sub-clusters (Supplementary Fig. S7F) that are described next. Ductal and lobular sub-clusters were identified by using the highest clustering resolution (1) (Fig. 5C). This resulted in 6 sub-clusters. The marker genes (Supplementary Table S11, Fig. 5C) for each sub-cluster were identified using Seurat’s “FindAllMarkers” function (top 5 markers shown in Supplementary Fig. S7G). To link scRNAseq sub-clusters with DSP ROI, all DSP ROIs were scored by sub-cluster marker genes using GSVA (Fig. 5D). Pathway analysis: hallmark pathway and biological theme analysis was performed as described above for DSP dataset.

### References

1.	Merritt, C. R. *et al.* Multiplex digital spatial profiling of proteins and RNA in fixed tissue. *Nature Biotechnology 2020 38:5* **38**, 586–599 (2020).

2.	Rueden, C. T. *et al.* ImageJ2: ImageJ for the next generation of scientific image data. *BMC Bioinformatics* **18**, 529 (2017).

3.	Team, R. C. R: A Language and Enviornment for Statistical Computing. Preprint at (2022).

4.	Krijthe, J. H. {Rtsne}: T-Distributed Stochastic Neighbor Embedding using Barnes-Hut Implementation. Preprint at (2015).

5.	van der Maaten, L. J. P. Accelerating t-SNE using Tree-Based Algorithms. *Journal of Machine Learning Research* **15**, 3221–3245 (2014).

6.	Gu, Z., Eils, R. & Schlesner, M. Complex heatmaps reveal patterns and correlations in multidimensional genomic data. *Bioinformatics* **32**, 2847–2849 (2016).

7.	Wilkerson, M. D. & Hayes, D. N. ConsensusClusterPlus: A class discovery tool with confidence assessments and item tracking. *Bioinformatics* **26**, 1572–1573 (2010).

8.	Haibe-Kains, B. *et al.* A Three-Gene Model to Robustly Identify Breast Cancer Molecular Subtypes. *JNCI: Journal of the National Cancer Institute* **104**, 311–325 (2012).

9.	Law, C. W., Chen, Y., Shi, W. & Smyth, G. K. Voom: Precision weights unlock linear model analysis tools for RNA-seq read counts. *Genome Biol* **15**, (2014).

10.	Ritchie, M. E. *et al.* limma powers differential expression analyses for RNA-sequencing and microarray studies. *Nucleic Acids Res* **43**, e47–e47 (2015).

11.	Ciriello, G. *et al.* Comprehensive molecular portraits of invasive lobular breast cancer. *Cell* **163**, 506–519 (2015).

12.	Rahman, M. *et al.* Alternative preprocessing of RNA-Sequencing data in The Cancer Genome Atlas leads to improved analysis results. *Bioinformatics* **31**, 3666–3672 (2015).

13.	Du, T. *et al.* Invasive lobular and ductal breast carcinoma differ in immune response, protein translation efficiency and metabolism. *Scientific Reports 2018 8:1* **8**, 1–11 (2018).

14.	Liberzon, A. *et al.* The Molecular Signatures Database Hallmark Gene Set Collection. *Cell Syst* **1**, 417–425 (2015).

15.	Kanehisa, M., Furumichi, M., Sato, Y., Kawashima, M. & Ishiguro-Watanabe, M. KEGG for taxonomy-based analysis of pathways and genomes. *Nucleic Acids Res* **51**, D587–D592 (2023).

16.	Liberzon, A. *et al.* Molecular signatures database (MSigDB) 3.0. *Bioinformatics* **27**, 1739–1740 (2011).

17.	Federico, A. & Monti, S. HypeR: An R package for geneset enrichment workflows. *Bioinformatics* **36**, 1307–1308 (2020).

18.	Hänzelmann, S., Castelo, R. & Guinney, J. GSVA: Gene set variation analysis for microarray and RNA-Seq data. *BMC Bioinformatics* **14**, 1–15 (2013).

19.	Ashburner, M. *et al.* Gene ontology: tool for the unification of biology. The Gene Ontology Consortium. *Nat Genet* **25**, 25–29 (2000).

20.	Carbon, S. *et al.* The Gene Ontology resource: enriching a GOld mine. *Nucleic Acids Res* **49**, D325–D334 (2021).

21.	Gillespie, M. *et al.* The reactome pathway knowledgebase 2022. *Nucleic Acids Res* **50**, D687–D692 (2022).

22.	da Silva, E. M. *et al.* Mesonephric and mesonephric-like carcinomas of the female genital tract: molecular characterization including cases with mixed histology and matched metastases. *Mod Pathol* **34**, 1570–1587 (2021).

23.	da Silva, E. M. *et al.* TERT promoter hotspot mutations and gene amplification in metaplastic breast cancer. *NPJ Breast Cancer* **7**, (2021).

24.	Selenica, P. *et al.* APOBEC mutagenesis, kataegis, chromothripsis in EGFR-mutant osimertinib-resistant lung adenocarcinomas. *Ann Oncol* **33**, 1284–1295 (2022).

25.	Da Cruz Paula, A. *et al.* Genomic profiling of primary and recurrent adult granulosa cell tumors of the ovary. *Mod Pathol* **33**, 1606–1617 (2020).

26.	da Silva, E. M. *et al.* Mesonephric and mesonephric-like carcinomas of the female genital tract: molecular characterization including cases with mixed histology and matched metastases. *Mod Pathol* **34**, 1570–1587 (2021).

27.	Zehir, A. *et al.* Mutational landscape of metastatic cancer revealed from prospective clinical sequencing of 10,000 patients. *Nat Med* **23**, 703–713 (2017).

28.	Cheng, D. T. *et al.* Memorial Sloan Kettering-Integrated Mutation Profiling of Actionable Cancer Targets (MSK-IMPACT): A Hybridization Capture-Based Next-Generation Sequencing Clinical Assay for Solid Tumor Molecular Oncology. *The Journal of Molecular Diagnostics* **17**, 251–264 (2015).

29.	Li, H. & Durbin, R. Fast and accurate short read alignment with Burrows-Wheeler transform. *Bioinformatics* (2009) doi:10.1093/bioinformatics/btp324.

30.	McKenna, A. *et al.* The Genome Analysis Toolkit: a MapReduce framework for analyzing next-generation DNA sequencing data. *Genome Res* **20**, 1297–1303 (2010).

31.	Cibulskis, K. *et al.* Sensitive detection of somatic point mutations in impure and heterogeneous cancer samples. *Nat Biotechnol* **31**, 213–219 (2013).

32.	Saunders, C. T. *et al.* Strelka: accurate somatic small-variant calling from sequenced tumor-normal sample pairs. *Bioinformatics* **28**, 1811–1817 (2012).

33.	Koboldt, D. C. *et al.* VarScan 2: somatic mutation and copy number alteration discovery in cancer by exome sequencing. *Genome Res* **22**, 568–576 (2012).

34.	Narzisi, G. *et al.* Accurate de novo and transmitted indel detection in exome-capture data using microassembly. *Nature Methods 2014 11:10* **11**, 1033–1036 (2014).

35.	Narzisi, G. *et al.* Genome-wide somatic variant calling using localized colored de Bruijn graphs. *Commun Biol* **1**, (2018).

36.	Chang, M. T. *et al.* Accelerating Discovery of Functional Mutant Alleles in Cancer. *Cancer Discov* **8**, 174–183 (2018).

37.	Li, H. *et al.* The Sequence Alignment/Map format and SAMtools. *Bioinformatics* **25**, 2078–2079 (2009).

38.	Shen, R. & Seshan, V. E. FACETS: allele-specific copy number and clonal heterogeneity analysis tool for high-throughput DNA sequencing. *Nucleic Acids Res* **44**, (2016).

39.	Carter, S. L. *et al.* Absolute quantification of somatic DNA alterations in human cancer. *Nat Biotechnol* **30**, 413–421 (2012).

40.	Gulhan, D. C., Lee, J. J. K., Melloni, G. E. M., Cortés-Ciriano, I. & Park, P. J. Detecting the mutational signature of homologous recombination deficiency in clinical samples. *Nat Genet* **51**, 912–919 (2019).

41.	Alexandrov, L. B. *et al.* Signatures of mutational processes in human cancer. *Nature* **500**, 415–421 (2013).

42.	Alexandrov, L. B. *et al.* The repertoire of mutational signatures in human cancer. *Nature* **578**, 94–101 (2020).

43.	Vahdatinia, M. *et al.* KIT genetic alterations in breast cancer. *J Clin Pathol* (2022) doi:10.1136/JCP-2022-208611.

44.	Geyer, F. C. *et al.* Genetic analysis of a morphologically heterogeneous ovarian endometrioid carcinoma. *Histopathology* **71**, 480 (2017).

45.	Hao, Y. *et al.* Integrated analysis of multimodal single-cell data. *Cell* (2021) doi:10.1016/j.cell.2021.04.048.

46.	Franzén, O., Gan, L. M. & Björkegren, J. L. M. PanglaoDB: a web server for exploration of mouse and human single-cell RNA sequencing data. *Database* **2019**, 46 (2019).

47.	Zhang, L. *et al.* Single-Cell Analyses Inform Mechanisms of Myeloid-Targeted Therapies in Colon Cancer. *Cell* **181**, 442-459.e29 (2020).

48.	Gao, R. *et al.* Delineating copy number and clonal substructure in human tumors from single-cell transcriptomes. *Nature Biotechnology 2021 39:5* **39**, 599–608 (2021).
